# Functional characterization of CCHamides and deorphanization of their receptors in the yellow fever mosquito, *Aedes aegypti*

**DOI:** 10.1101/2024.01.27.577569

**Authors:** Jinghan Tan, Susanne Neupert, Jean-Paul Paluzzi

## Abstract

As a widely distributed anthropophilic mosquito species and vector of various arboviruses, *Aedes aegypti* poses a significant threat to human health on a global scale. Investigating mosquito neuropeptides allows us to better understand their physiology. The neuropeptides CCHamide1 (CCHa-1) and CCHamide-2 (CCHa-2) and their associated G protein-coupled receptors (CCHa-1R and CCHa-2R) were recently identified and studied across insects. However, expression profiles and physiological roles of CCHamides and their receptors in many other insects, including *A. aegypti*, remain unclear. This research aimed to quantify and localize the expression of CCHamides along with their receptors and to elucidate their physiological function in the yellow fever mosquito. RT-qPCR analysis revealed transcript abundance of CCHamides and receptors changes over development. Differential expression was also observed in tissues/organs of adult mosquitoes indicating *CCHa-1* and *CCHa-2* transcripts are most highly enriched in the midgut, while receptors are expressed across various tissues. Further, CCHamides were immunolocalized in neurons in the brain and ventral nerve cord along with enteroendocrine cells in the posterior midgut adjacent to the midgut-hindgut junction, corroborating their transcript expression profiles. Using different mass spectrometrical approaches, presence of CCHamides were confirmed in the brain of both sexes, including the *pars intercerebralis* of female mosquitoes, as well as in the gut of adult mosquitoes. For chemical identification of predicted CCHamides, we analyzed brain and gut extracts by ESI-Q Exactive Orbitrap MS and resulting fragmentations confirmed CCHa1 and CCHa2 in brain and midgut samples of both male and female mosquitoes. A heterologous functional assay was used to confirm the specificity and sensitivity of the two CCHamide receptors by assessing their activation in response to diverse mosquito peptides, which confirmed CCHa-1 and CCHa-2 as the natural ligands. Finally, using a capillary feeder (CAFE) bioassay, our results suggest that CCHa-2 modulates feeding behaviour in female mosquitoes.

## 1. Introduction

The proper operation of the organism as a whole is inseparable from intercellular communication including signaling between the CNS and peripheral organs, which involves a wide range of signaling molecules. Neuropeptides, which are small proteinaceous compounds that normally consist of between 4 to 45 amino acid residues, are recognized as one of the most diverse and versatile groups of signaling molecules that can function as neurotransmitters acting at synapses, neuromodulators that regulate and affect the activity of other neurotransmission processes, and neurohormones that act on cells locally or at a distance (Agrawal et al., 2019; Burbach, 2011; Predel et al., 2010). Neurosecretory cells and endocrine cells are responsible for producing and releasing these small proteinaceous substances into the haemolymph of insects (as well as other animals) which then activate their cognate receptors on cells of target tissues to trigger signal transduction and downstream responses (Abou El Asrar et al., 2020).

Neuropeptides have a variety of primary structures and carry out numerous distinct functions. Well over 50 neuropeptides have been isolated and characterized over the years in the yellow fever mosquito, *Ae. aegypti* (Brown et al., 2008; Champagne and Ribeiro, 1994; Li et al., 2006; Matsumoto et al., 1989; Predel et al., 2010; Riehle et al., 2006; Stracker et al., 2002; Strand et al., 2016; Veenstra et al., 1997). A unique subset of gut-produced neuropeptides that are also enriched in the nervous system are classified as brain-gut peptides; through the brain-gut axis, not only can they act locally but can target other tissues at a distance. Studies have shown that they are important in modulating growth and development, reproduction, excretion and addiction behaviour across insects (Abou El Asrar et al., 2020; Chung et al., 1999; McBrayer et al., 2007; Pérez-Hedo et al., 2010; Vadnie et al., 2014; Vanderveken and O’Donnell, 2014; Wu et al., 2020).

One such family of brain-gut neuropeptides, CCHamides, were originally noted by Zdárek *et al*. in 2000 in their search for the parturition hormone in the tsetse fly, *Glossina morsitans*. CCHamides have since been discovered in arthropods, such as crustaceans, centipedes and insects, and subsequently chemically isolated, sequenced and functionally characterized in *D. melanogaster* (Chipman et al., 2014; Christie, 2014; Ida et al., 2012; Ren et al., 2015; Veenstra and Ida, 2014). Two separate genes were identified in insects that code for CCHamides, *CCHa1* (CG14358) and *CCHa2* (CG14375). Even though their amino acid sequences are different, both neuropeptides in *Drosophila* are structurally similar with a length of 13 amino acid residues, an amide functional group at the C-terminus and a disulfide bond formed by two cystines, which allows CCHamides to form a cyclical structure. Additionally, the *D. melanogaster* CCHamides also share a YGH motif and a C-terminal GXG-NH_2_ motif, where X denotes either an alanine or glycine residue (Ida et al., 2012).

The large family of membrane proteins known as G protein-coupled receptors (GPCRs) are the most common neuropeptide receptors and can initiate a signaling cascade involving either Ca^2+^ flux or other second messengers including IP_3_, cAMP and diacylglycerol (Abou El Asrar et al., 2020; Agrawal et al., 2019; Caers et al., 2012; Gilbert et al., 1988). Earlier studies established two *Drosophila* GPCRs, CCHa1R (CG14593) and CCHa2R (CG30106) as authentic receptors of the endogenous CCHa1 and CCHa2 ligands, respectively, demonstrating the highest specificity and sensitivity in activating these receptors (Hansen et al., 2011; Ida et al., 2012). Phylogenetic evidence suggested that CCHamide receptors belong to the bombesin/gastrin-releasing peptide receptor superfamily, which are sub-divided into gastrin-releasing peptide-preferring receptor (GRPR), the neuromedin B-preferring receptor (NMBR) and bombesin receptor subtype 3 (BRS3). Based on these phylogenetic analyses, CCHamide receptors share evolutionary relationship with the vertebrate bombesin/gastrin-releasing peptide receptor superfamily (Hansen et al., 2011; Ida et al., 2012; Ohki-Hamazaki et al., 1997; Tang et al., 2019; Veenstra and Ida, 2014).

Previous studies reported that mRNA transcript levels of CCHamides and CCHamide receptors fluctuate throughout developmental stages in *Drosophila*, exhibiting the highest enrichment in adults of both sexes. Notably, sexually dimorphic expression in adult flies of these neuropeptides and their receptors was also observed (Li et al., 2013). Differential expression patterns were found as *CCHa1* and *CCHa1R* were enriched in the brain and midgut region, but *CCHa2* was particularly enriched in the midgut, while the brain was determined as a nearly exclusive site expressing *CCHa2R* (Li et al., 2013). These expression profiles might therefore suggest that CCHa1 and CCHa2 hold distinct physiological roles through different signaling pathways despite possible cross-activation of their receptors as observed at higher concentrations (Hansen et al., 2011; Ida et al., 2012). As one example, both neuropeptides were linked to feeding to some extent since lack of CCHa1 or CCHa2 resulted in a reduction in foraging activity (Farhan et al., 2013; Ren et al., 2015). However, further studies demonstrated that a protein-rich meal led to increased expression of *CCHa1* in *Drosophila*, while CCHa2 acted as a sugar-responsive neuropeptide (Ren et al., 2015; Titos and Rogulja, 2020). In most cases, changes in expression of *CCHa1* and *CCHa2* that are induced by feeding affect other physiological processes. Loss of CCHa1 led to a disruption to normal circadian rhythm and higher arousability in *Drosophila*, while CCHa2 seems to play a role in development since pupariation time was delayed by nearly three days due to the lack of functional *CCHa2* transcript (Fujiwara et al., 2018; Ren et al., 2015; Titos and Rogulja, 2020). In the pea aphid, *Acyrthosiphon pisum*, the expression of the presumed CCHa2 receptor was found to be significantly increased in starved aphids while knockdown by RNAi led to a reduction in food intake and reproductive output but had no influence on survival (Shahid et al., 2021). Recent work in the two-spotted cricket, *Gryllus bimaculatus*, revealed that synthetic CCHa2 (GbCCHa-2) injection reduced food intake (as determined by the number of fecal pellets excreted) and increased carbohydrate and lipid levels in the haemolymph while knockdown of its receptor (GbCCHa-2R) increased food intake and led to a corresponding reduction in haemolymph lipid levels (Zhu et al., 2022).

As a widely distributed anthropophilic mosquito species, *Ae. aegypti* transmits various human disease agents. Rapid urbanization and global warming have allowed these mosquitoes to invade more areas around the world escalating the risks of infectious diseases to human communities (Jansen and Beebe, 2010; Reinhold et al., 2018). Studying CCHamides and their receptors allows us to better understand mosquito biology and could help in preventing mosquito-borne diseases through new strategies for vector control based on insight on these neuropeptides. This study aimed to characterize CCHamides in *Ae. aegypti* via mass spectrometry and immunohistochemistry, quantify mRNA expression of CCHamides and their receptors and deorphanize both receptors to help unravel their physiological functions in the yellow fever mosquito as this information remains unknown.

## 2. Materials and Methods

### 2.1 Animal rearing

*Ae. aegypti* (Liverpool strain) colony maintenance involved regular blood feeding of female mosquitoes with sheep’s blood in Alsever’s solution (Cedarlane Laboratories, Burlington, ON), which allows them to produce eggs. Dampened Whatman filter papers (GE Bioscience) were placed in cages housing adult *Ae. aegypti* providing a suitable substrate for egg laying by females. The eggs are collected, stored semi-desiccated for up to several months and are then placed in a plastic container with distilled water to hatch larvae reared under a 12:12 hr photoperiod in an incubator at 25°C as previously described (Wahedi and Paluzzi, 2018). Hatched larvae were fed once every two days with several drops of liquid larval food comprised of 2% (w/v) beef liver powder (NOW Foods, Bloomingdale, Illinois) and 2% (w/v) brewer’s yeast (Bulk Barn, Milton, ON) in distilled water. Unless noted otherwise, all adult mosquitoes were provided with a 10% sucrose solution in a small tube fitted with a cotton ball. Pupae were isolated and transferred into small plastic mesh-covered containers (BudDorm, MegaView Science, Taiwan) and the resulting adult mosquitoes were either used for subsequent experiments or transferred back to the main colony in large mesh covered plastic containers (BudDorm, MegaView Science, Taiwan).

### 2.2 CCHamide and receptor expression profiles

Whole mosquitoes of different developmental stages and dissected tissues of one-day-old adult male and female mosquitoes (n=35-40) were mechanically homogenized with a disposable polypropylene pestle. Total RNA was then extracted using a Monarch® Total RNA Miniprep Kit (New England Biolabs, Whitby, ON, Canada) following the manufacturer’s protocol. Whole insect total RNA (500ng) or tissue-specific total RNA (80ng) was used as a template to synthesize cDNA using iScript^TM^ Reverse Transcription Supermix (Bio-Rad, Mississauga, ON) followed by an 11-fold dilution with nuclease-free water prior to RT-qPCR analysis.

Expression profiles of *CCHa1*, *CCHa2, CCHa1R* and *CCHa2R* were assessed by amplifying a 164bp, 227bp, 74bp and 226bp DNA fragment, respectively, with optimized primers designed using Geneious Pro Bioinformatics Software (Supplement Table S1) on a StepOnePlus Real-Time PCR System (Applied Biosystems, Carlsbad, CA, United States) using PowerUP^TM^ SYBR® Green Master Mix (Applied Biosystems, Carlsbad, CA, United States). The correctness of amplicons was confirmed by Sanger sequencing (Center for Applied, Genomics, Hospital for Sick Children, Toronto, ON) and compared to the reference genome sequence of *Ae. aegypti CCHa1* (GenBank accession: XM_021854293.1), *CCHa1R* (GenBank accession: XM_021851237.1), *CCHa2* (GenBank accession: XM_021838098.1) and *CCHa2R* (GenBank accession: XM_021852958.1). RT-qPCR cycling condition started at 50°C for 2 minutes followed by 95°C for 20 seconds, and then 40 cycles of (i) 95°C for 3 seconds and (ii) 62°C for 30 seconds. Two reference genes, ribosomal protein 49 (*rp49*) and *rpS18*, were used as optimal endogenous controls to normalize the expression of target genes of interest via ΔΔC_T_ method using primers described previously (Paluzzi et al., 2014). Quantitative determination of expression profiles of the two *Aedae*CCHamide transcripts*, CCHa1R* and *CCHa2R* were analyzed in 3-5 biological replicates with each containing triplicate technical replicates and no-template controls.

### 2.3 Enzyme-linked immunosorbent assay (ELISA)

A custom rabbit polyclonal affinity-purified antibody, anti-*Aedae*CCHa2 (antigen sequence: Cys-GCAAFGHACYGGH-NH_2_) (Biomatic Co., Kitchener, ON), was used to target *Aedae*CCHa2 neuropeptide in mosquito tissues via whole mount immunohistochemistry. An ELISA test was conducted to determine the optimal antibody concentration, examine the binding specificity to the CCHa2 neuropeptide and identify potential cross-reactivity with other neuropeptides, including CCHa1.

Serial dilutions of synthetic *Aedae*CCHa1, *Aedae*CCHa2, *Aedae*RYamide1 (Ida et al., 2011) and *Aedae*CAPA-PK1 (Predel et al., 2010) (PepMic, Suzhou, China) (Supplement Table S2) were prepared ranging from 0.05μmol to 10μmol in 100μL carbonate buffer that consisted of 15mmol/L Na_2_CO_3_-H_2_O (pH 9.4) and 35mmol/L NaHCO_3_ (pH 9.4), and loaded onto a 96-well high binding ELISA plate (Sarstedt Inc. Montreal, QC) covered with plastic wrap, and then incubated at 4°C for 48h. The contents of the wells were then emptied and rinsed three times using wash solution (346mmol/L NaCl, 2.7mmol/L KCl, 1.5mmol/L KH_2_PO_4_, 5.1mmol/L Na_2_HPO_4_-7H_2_O and 0.5% Tween-20). Wells were then treated with anti-*Aedae*CCHa2 primary antibody (1:1000 dilution) prepared in block solution containing 0.5% (w/v) skimmed milk powder and protease-free bovine serum albumin (BSA) made up in DPBS and incubated for 2h at room temperature on a shaking platform. Following primary antibody incubation, each well underwent three washes before the application of a secondary antibody (1:1500 dilution), anti-rabbit-HRP (Bio-Rad Laboratories (Canada) Ltd, Mississauga, ON) and incubated for 1h at room temperature on a rocker. This was followed by three washes and then tetramethylbenzidine (TMB) solution (Sigma-Aldrich Canada Co., Oakville, ON) was loaded into each well of the plate, and 2N HCl solution was added once the colour had developed. A Synergy 2 Multi-Mode Microplate Reader (BioTek, Winooski, VT, USA) was used to measure the absorbance of each well at 450 nm.

### 2.4 Whole mount immunohistochemistry

In order to localize the distribution of CCHamide-producing neurons and endocrine cells, and in light of the examined transcript expression profile, the entire gut region, including the midgut, pyloric valve (i.e. the junction between the midgut and hindgut), the hindgut, along with the brain and abdominal ganglia of adult male and female sugar-fed mosquitoes, *CCHa2* knockdown mosquitoes and 24 to 48h starved mosquitoes were dissected to perform whole mount immunohistochemistry, following a protocol described previously (Sajadi et al., 2020). Dissected tissues were fixed in 4% paraformaldehyde (PFA) overnight at 4°C. Tissues were washed three times for 10 minutes with DPBS and then incubated for 1h at room temperature with 4% Triton X-100, 2% bovine serum albumin (BSA) and 10% normal sheep serum (NSS) for tissue permeabilization. Primary antibody (5μg/mL), anti-*Aedae*CCHa2 (Biomatic Co., Kitchener, ON) described above, was prepared in antibody dilution buffer containing 0.4% Triton X-100 (v/v), 2% NSS, 2% BSA, Triton X-100 in DPBS 2h prior to use, and then incubated with permeabilized tissues for 48h at 4°C on a rocking platform with gentle agitation. For control treatments, 10μM of either CCHa1 or CCHa2 synthetic peptide (PepMic, Suzhou, China) was pre-incubated overnight along with the primary antibody and used on permeabilized tissues as described above.

After primary antibody incubation, tissues were washed three times with DPBS and then incubated for 24h at 4°C on a rocking platform with the secondary antibody, goat anti-rabbit Alexa 568 (Life Technologies, Burlington, ON) that was diluted 1:200 in DPBS containing 10% NSS. Tissue samples were subsequently rinsed three times with DPBS again and then transferred onto microscope slides in mounting buffer containing 1:1 DPBS: glycerol and 1µg/mL 4’,6-diamidino-2-phenylindole dihydrochloride (DAPI) (Molecular Probes, Eugene, OR) to stain cell nuclei in prepared tissues and examined under a Lumen Dynamics XCite™ 120Q Nikon fluorescence microscope (Nikon, Mississauga, ON, Canada).

### 2.5 Tissue preparation for mass spectrometry

*Ae. aegypti* were anesthetized on ice. Head capsules and the abdomen were opened and covered with ice-cold physiological insect saline (128 mM NaCl, 2.7 mM KCl, 2 mM CaCl_2_, 1.2 mM NaHCO_3_, pH 7.25). Direct tissue profiling (n=15) involved brain and digestive system being removed and transferred into a drop of fresh saline. Small parts of different brain areas including *pars intercerebralis* as well as the gut system subdivided into esophagus and posterior midgut, were separately dissected and transferred using a glass capillary fitted to a tube with a mouthpiece into a drop of water placed on a sample plate for MALDI-TOF MS analysis as described in a previous study (Schachtner et al., 2010). Immediately after transfer, water was removed around the sample and the tissue was allowed to dry before matrix application.

*Tissue extraction (in total n=6)*: We made six different sample sets, consisting of 20, 30 and 50 heads/brains, and separately 20, 30 and 50 midguts. Each sample set was collected in 30μL extraction solution containing 50 % methanol, 49 % H_2_O and 1 % formic acid (FA) on ice. Tissue samples were homogenized using an ultrasonic bath (Transonic 660/H, Elma Schmidbauer GmbH, Hechingen, Germany) for 90 minutes on ice. Afterwards, the samples were centrifuged for 15 minutes at 13,000 rpm at 4 °C. The supernatants were separated and then evaporated in a vacuum concentrator to remove organic solvent. Extracts were stored at –20 °C until use.

### 2.6 Matrix application

*Direct tissue profiling:* 10 mg/mL 2,5–dihydroxybenzoic acid (DHB, Sigma-Aldrich, Steinheim, Germany) dissolved in 20 % acetonitrile/1 % FA/79 % water (Fluka). Dried tissue samples were covered with 0.1 µL DHB solution applied by a 0.1-2.5 µL Eppendorf pipette (Eppendorf AB, Hamburg, Germany). For extract analysis, 0.1 µL concentrated supernatant was mixed with 0.1 µL DHB solution by a 0.1-2.5 µL Eppendorf pipette. Subsequently, sample spots were blow-dried with a commercially available hair dryer to form homogeneous crystals.

### 2.7 MALDI-TOF MS

Mass spectra were acquired manually using an Rapiflex TOF/TOF mass spectrometer (Bruker Daltonik GmbH, Bremen, Germany) in reflector positive ion mode in a mass range of 600–4,000 Da. For calibration, the following synthetic peptide mixture was used: proctolin, *Drosophila melanogaster* (Drm)-sNPF-1^4-11^, *Locusta migratoria* periviscerokinin-1 (PVK-1), *Periplaneta americana* (Pea)-FMRFa-12, *Manduca sexta* allatotropin (AT), Drm-IPNa, Pea-SKN and glucagon. Laser fluency was adjusted to provide an optimal signal-to-noise ratio. The obtained data were processed using FlexAnalysis V.3.4 software package.

### 2.8 Quadrupole Orbitrap MS Coupled to Nanoflow HPLC

Before injecting the samples into the nanoLC system, extracts were desalted using self-packed Stage Tip C18 (IVA Analysentechnik e.K., Meerbusch, Germany) spin columns (Rappsilber et al., 2007). For analysis, peptides were separated on an EASY nanoLC 1000 UPLC system (Thermo Fisher Scientific, Waltham, MA) using RPC18 columns 50 cm (fused Silica tube with ID 50μm ± 3μm, OD 150μm ± 6μm, Reprosil 1.9μm, pore diameter 60 Å, Dr. Maisch, Ammerbuch-Entringen, Germany) and a binary buffer system (A: 0.1 % FA, B: 80 % ACN, 0.1 % FA) as described for *Cataglyphis nodus* samples (Habenstein et al., 2021). Running conditions were as follows: linear gradient from 2 to 62 % B in 110 min, 62 to 75 % B in 30 min, and final washing from 75 to 95 % B in 6 min (45 °C, flow rate 250nL/min). Finally, the gradients were re-equilibrated for 4 min at 5 % B. The HPLC was coupled to a Q-Exactive Plus (Thermo Scientific, Bremen, Germany) mass spectrometer. MS data were acquired in a top-10 data-dependent method dynamically choosing the most abundant peptide ions from the respective survey scans in a mass range of 300−3,000 m/z for HCD fragmentation. Full MS^1^ acquisitions ran with 70,000 resolution, with automatic gain control target (AGC target) at 3e6 and maximum injection time at 80ms. HCD spectra were measured with a resolution of 35,000, AGC target at 3e6, maximum injection time at 240ms, 28 eV normalized collision energy, and dynamic exclusion set at 25 s. The instrument was run in peptide recognition mode (i.e., from two to eight charges), singly charged and unassigned precursor ions were excluded. Raw data were analyzed with PEAKS Studio 10.5 (Bioinformatics Solutions Inc., ON, Canada). Neuropeptides were searched against an internal database comprising *Ae. aegypti* neuropeptide precursor sequences with parent mass error tolerance of 0.2 Da and fragment mass error tolerance of 0.2 Da. Setting enzymes: none was selected because samples were not digested. The false discovery rate (FDR) was enabled by a decoy database search as implemented in PEAKS 10.5. The following posttranslational modifications (PTM) were selected: C-terminal amidation and half of disulfide bridges as fixed PTM, and oxidation at methionine, phosphylation and sulfatation as variable PTMs. In each run, a maximum of three variable PTMs per peptide were allowed. Fragment spectra with a peptide score (−10 lgP) equivalent to a P-value of ∼1 %, were manually reviewed.

### 2.9 Heterologous functional activity bioluminescence assays

The complete open reading frames of *Aedae*CCHa1R and *Aedae*CCHa2R were amplified by PCR using Q5 high-fidelity DNA polymerase (New England Biolabs, Whitby, ON, Canada) with primers located upstream of the start codon and downstream of the stop codon (Supplement Table S1) based on the *Ae. aegypti* predicted receptor sequences (CCHa1R GenBank accession: XM_021851237.1; CCHa2R GenBank accession: XM_021852958.1). Following this initial amplification, PCR products were purified with a Monarch PCR & DNA clean-up kit (New England Biolabs, Whitby, ON, Canada) and used as template to reamplify the products to incorporate a Kozak translation initiation sequence prior to the start codon (Kozak, 1986) along with restriction enzymes for directional cloning following standard protocols described previously (Gondalia et al., 2016; Oryan et al., 2018; Wahedi and Paluzzi, 2018). These amplified products were then purified as above and restriction enzyme digested CCHa1R and CCHa2R sequences were ligated into pcDNA3.1+ and pBudCE4.1 mammalian expression vectors followed by bacterial transformation and plasmid DNA purification to obtain a large quantity of receptor constructs using ZymoPURE II Plasmid Midiprep Kit (Zymo Research, Tustin, CA, USA) following the manufacturer recommendation. Base-pair accuracy of mammalian expression constructs was confirmed by Sanger sequencing (Center for Applied, Genomics, Hospital for Sick Children, Toronto, ON).

A recombinant Chinese hamster ovary cell (CHO-K1) line stably expressing aequorin (Gondalia et al., 2016; Paluzzi et al., 2012) was used for transiently expressing *Aedae*CCHa1R and *Aedae*CCHa2R. Cell growth and colony maintenance followed the optimized protocol as previously reported (Wahedi and Paluzzi, 2018). Freshly seeded cells between 90% to 100% confluency were used for transfection. Cells were transfected with either pcDNA3.1+ receptor construct, pBudCE4.1 receptor construct containing the murine promiscuous G_alpha_15 (Paluzzi et al., 2014) or pBud-EGFP, using Lipofectamine 3000 or LTX reagent (Invitrogen, Burlington, ON) following the manufacture guidelines. Cells transfected with the plasmid constructs were expected to produce and express either CCHa1R, CCHa2R or green fluorescent protein (EGFP) with the latter acting as a positive control for transfection and negative control for the bioluminescence receptor functional assay.

Cells were harvested 48h post-transfection by dislodging with DPBS-EDTA solution, centrifugating and resuspending in assay media (BSA media), which was prepared with DMEM: F12 media, 10% Bovine Serum Albumin (BSA) and Antibiotic-Antimycotic, with 5μM coelenterazine-h. Following a 3h incubation at room temperature with stirring in the dark, cells were further diluted 10-fold in assay media and incubated for an additional 45 minutes with 0.5μM coelenterazine-h final concentration following dilution. In order to examine the selectivity of responsiveness of *Aedae*CCHa1R and *Aedae*CCHa2R to multiple neuropeptides, a library of commercially synthesized insect neuropeptides (Supplement Table S2) at a purity of >90% were prepared (10^-6^M and 10^-7^M final titre) in assay media and loaded on 96-well luminescence plates (Greiner Bio-One, Germany) in quadruplicates following a palindromic loading scheme. To generate dose-response curves of *Aedae*CCHa1 and *Aedae*CCHa2 to both receptors, serial dilutions with concentrations ranging from 10^-5^M to 10^-14^M were prepared in BSA media and loaded on 96-well luminescence plates. BSA media alone was used to determine any background and acted as a negative control; 50μM ATP was used as a positive control to normalize the bioluminescent response of each well due to Ca^2+^ flux triggered by endogenous purinoceptors expressed in CHO-K1 cells (Iredale and Hill, 1993). Cells prepared for the functional assay were loaded into each well of a white 96-well plate using an automated injector unit, and the luminescent response was measured for 20 seconds followed by additional measurements for 10 seconds after ATP injection to the same well with a Synergy 2 Multi-Mode Microplate Reader (BioTek, Winooski, VT, USA).

### 2.10 Starvation assay

Adult mosquitoes of both sexes between 0 to 6h post-eclosion were isolated and randomly divided into containers with different diet conditions to examine the effects of starvation stress on the transcript abundance of *CCHa2* and *CCHa2R*. Test conditions included 10% sucrose-fed, water-fed and the combined starved/water-deprived for 24h. Mosquitoes were then collected and dissected for whole mount immunohistochemistry or used for RT-qPCR analysis.

### 2.11 Blood feeding assay

Hatched mosquito females were reared under normal laboratory conditions with males as previously described until reaching the age of 5-7 day-old without a single blood meal. They were then provided with pre-warmed (∼37°C) sheep blood (Cedarlane Laboratories, Burlington, ON) for 20 minutes using an artificial feeding membrane as described previously (Rocco et al., 2017). Blood-fed females, which can easily be distinguished by the red colour of their ballooned abdomen, were isolated and kept in containers with 10% sucrose solution for 1h, 6h, 12h and 24h post-blood feeding before whole body total RNA was isolated for subsequent RT-qPCR analysis to examine expression profiles of *CCHa1* and *CCHa2* in association with blood-feeding.

### 2.12 RNA interference (RNAi)

Two separate double-stranded RNA (dsRNA) target cDNA sequences for *A. aegypti CCHa2* transcript were amplified using gene-specific primers (Supplement Table S1). The target regions of these dsRNA span either partially within the 5’ untranslated region (UTR) and 5’end of the open reading frame or entirely within the 3’ UTR, with neither overlapping areas targeted by qPCR primers and thus avoiding false positive detection of *CCHa2* mRNA transcript. The dsRNA cDNA template for the EGFP gene was also amplified and used as a negative control. The dsRNA cDNA templates were ligated into the pGEM T-Easy vector (Promega, Madison, WI, United States) and subcloned into the dsRNA-producing vector L4440 (Fire et al., 1998; Montgomery et al., 1998). Purified dsRNA templates were reamplified from the L4440 vector with T7 promoter oligonucleotide sequences and used for dsRNA synthesis by *in vitro* transcription using HiScribe^®^ T7 High Yield RNA Synthesis Kit (New England Biolabs, Whitby, ON, Canada) following the manufacturer’s protocol. dsRNA denaturation and rehybridization, which helps to reduce secondary structures of RNA, were achieved by 5min incubation at 75°C and 15min incubation at room temperature, respectively. The Monarch^®^ RNA purification kit (New England Biolabs, Whitby, ON, Canada) was used to purify synthesized dsRNA following the manufacturer’s recommendation. The eluted dsRNA was concentrated to approximately 5μg/μL via vacuum centrifugation.

One-day-old adult mosquitoes were then injected in the thorax with 1μg (200nL) synthesized CCHa2 or EGFP dsRNA using a Nanoject III Programmable Nanoliter Injector (Drummond Scientific, Broomall, PA, USA). The knockdown efficiency of *CCHa2* expression in 4-day-old female mosquitoes was evaluated by RT-qPCR.

### 2.13 Capillary feeder assay (CAFE)

*CCHa2* transcript abundance and the circulating CCHa2 neuropeptide in one-day-old female mosquitoes were transiently knocked down or elevated by direct injection of dsRNA or commercially synthesized *Aedae*CCHa2 (10pmol dissolved in DPBS), respectively. Post-injected mosquitoes were allowed to recover for 2 days with a 10% sucrose solution provided prior to an 18 to 20h-starvation period. Subsequently, individual mosquitoes were transferred to small glass vials with rubber lids that allowed a capillary tube (VWR, Mississauga, ON, Canada) filled with 5μL of 10% sucrose to be inserted upright into each vial. The animals were incubated at 25°C for 24h. The amount of sucrose ingested by non-injected control, ds*EGFP*-injected, ds*CCHa2*-injected (*CCHa2* knockdown) and *Aedae*CCHa2-injected (CCHa2 elevated) mosquitoes was measured. Additionally, the amount of sucrose evaporated over 24h was recorded by inserting a capillary tube into an empty vial, and this value was subtracted from measured experimental values to account for background rates of evaporation.

### 2.14 Statistical analysis

All log-transformed RT-qPCR data and luminescence responses of multiple ligands in the heterologous functional assay and CAFE assay were analyzed by one-way ANOVA with Bonferroni’s multiple comparisons test and student’s t-test where p < 0.05 was considered significant. Dose-response curves and EC_50_ values were generated using the sigmoidal dose-response model. Statistical analysis was carried out by GraphPad Prism 9 (GraphPad Software, San Diego, USA).

## 3. Results

### 3.1 Developmental expression profiles

Transcript abundance of *CCHa1*, *CCHa2*, *CCHa1R* and *CCHa2R* demonstrated a dynamic profile over the course of post-embryonic mosquito development (Fig.1). In general, both neuropeptide transcripts showed a gradually increasing trend from pupal stage to adulthood despite that in the 4^th^ instar larval stage, *CCHa1* had higher transcript abundance (Fig.1A) while *CCHa2* had very low abundance (Fig.1C). Moreover, the highest *CCHa2* expression was detected in the early adulthood of both sexes (Fig.1C), but *CCHa1* transcript was enriched in 4-day old males (Fig.1A). Noticeably, differential expression of both neuropeptide transcripts between sexes of 4-day old adult mosquitoes was observed where males had significantly higher *CCHa1* and *CCHa2* transcript abundance compared to age-matched females. With respect to the two CCHamide receptor transcript profiles, *CCHa1R* transcript expression shared a similar expression profile to *CCHa1* with 4-day-old males having the highest abundance (Fig. 1B). In contrast, *CCHa2R* transcript expression was found to be more stable with similar abundance throughout developmental stages examined (Fig.1D). Even though the highest expression of *CCHa2R* was detected in the late pupal stage, it was not significantly different from most other stages, except for the 4-day old female mosquitoes.

**Figure 1.**
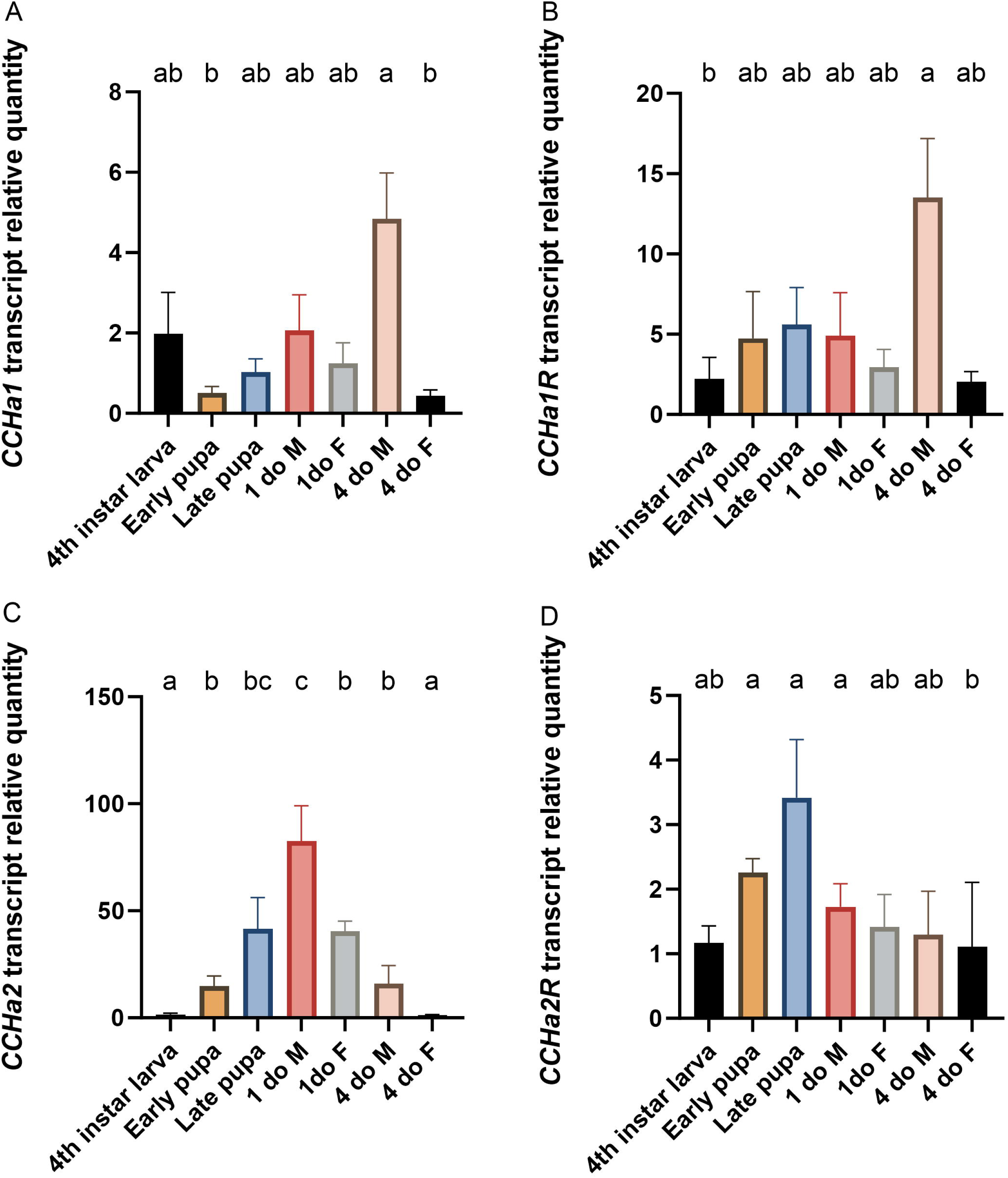
Expression profile of *Ae. aegypti CCHa1* (A), *CCHa1R* (B), *CCHa2* (C) and *CCHa2R* (D) in different post-embryonic developmental stages as determined by RT-qPCR. Abbreviations: 1do M = 1 day old male adult, 1do F = 1 day old female adult, 4do M = 4 day old male adult, 4do F = 4 day old female adult. Statistical differences are denoted with different letters, as determined by a one-way ANOVA on log-transformed data (p<0.05). Data represents the mean ± SEM (n=6).

### 3.2 Spatial expression profiles

Tissue and organ-specific expression patterns of CCHamide transcripts (*CCHa1* and *CCHa2*) and their receptors (*CCHa1R* and *CCHa2R*) were examined in 1-day-old male and female mosquitoes.

*CCHa1* transcript was distributed mainly in three tissues including the head (comprised mainly of the brain), midgut and carcass containing skeletal muscle, fat body and cuticle (Fig.2A-B). The midgut was determined to have a highly enriched *CCHa1* transcript in both sexes; however, differential expression between sexes was observed in other tissues. The abundance of *CCHa1* in the head of male mosquitoes was lower than the midgut region (Fig.2A) but had a similar abundance in the female head and midgut (Fig.2B). Even though the carcass was also enriched with *CCHa1* transcript compared to Malpighian tubules (MTs), its abundance was only a fraction of the levels detected in midgut of both males and females.

**Figure 2.**
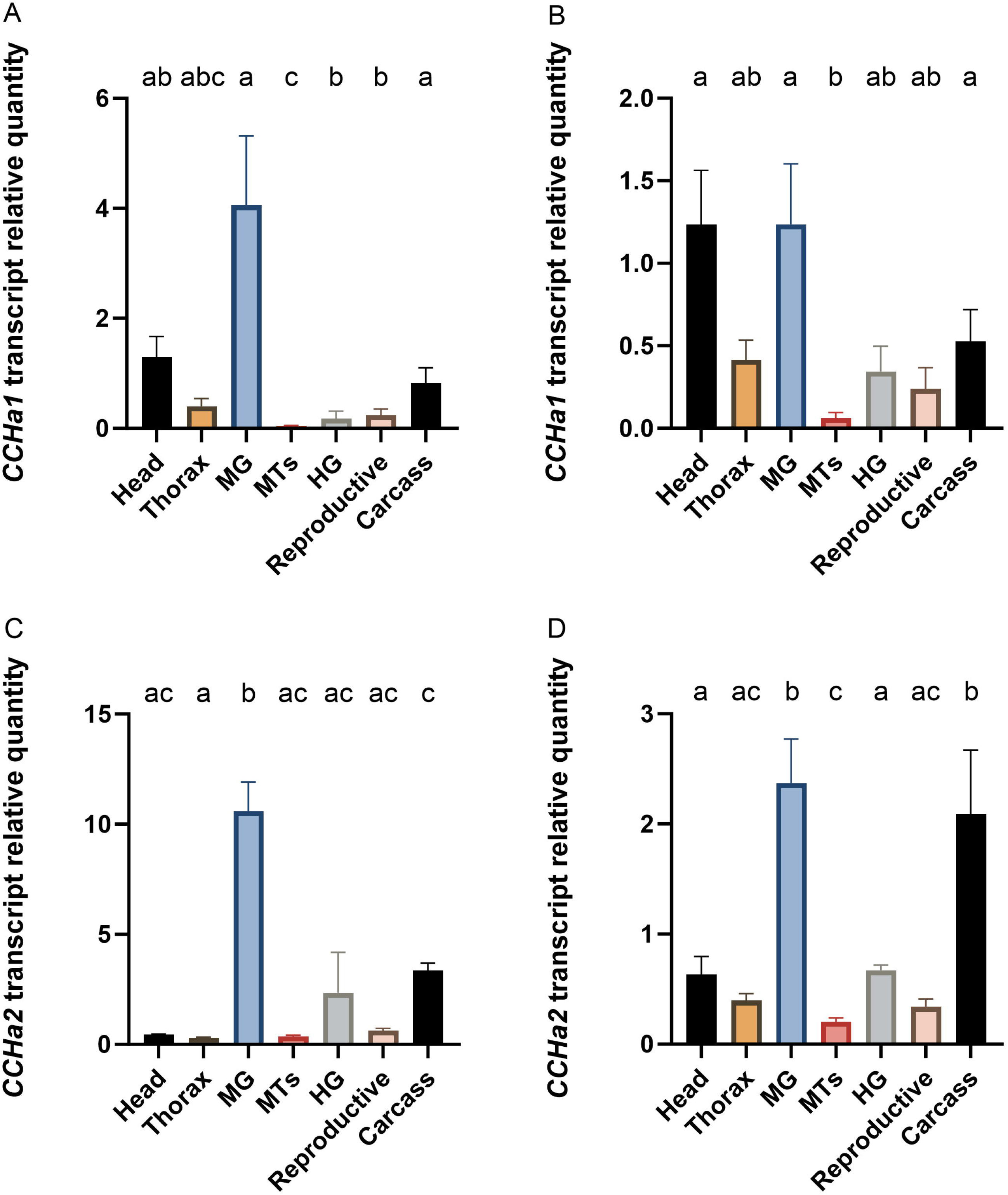
Tissue-specific expression profile of *CCHa1* in male (A) and female (B) along with *CCHa2* in male (C) and female (D) 1-day-old adult *Ae. aegypti* determined by RT-qPCR. Tissue/organ abbreviations: head, thorax, midgut (MG), Malpighian tubules (MTs), hindgut (HG), reproductive system and carcass. Statistical differences are denoted with different letters, as determined by a one-way ANOVA on log-transformed data (p<0.05). Data represents the mean ± SEM (n=4).

Similar to *CCHa1* expression, *CCHa2* transcript was significantly higher in midgut compared to all other examined tissues except for the female carcass, which also had a high abundance of *CCHa2* transcript (Fig.2C-D). Unlike the high *CCHa1* transcript levels in the head, the *CCHa2* expression was low.

Compared to the peptide transcripts that demonstrated enrichment in a subset of the examined tissues, the CCHa1 and CCHa2 receptor transcripts (*CCHa1R* and *CCHa2R*) were enriched in a different subset of tissues with differential expression between male and female mosquitoes (Fig.3). Unlike the observed *CCHa1* and *CCHa2* enrichment in the midgut, transcripts of CCHamide receptors were not found to be enriched in this tissue. Instead, the hindgut tended to be the site that demonstrated high *CCHa1R* abundance (Fig.3A-B). In addition, the head and carcass of male mosquitoes also demonstrated *CCHa1R* transcript enrichment (Fig.3A). *CCHa2R* transcript was found to be enriched in the male hindgut followed by the reproductive organs, the head and carcass (Fig.3C). In females, while there was no significant enrichment of *CCHa2R* in any examined tissue, the data suggests higher *CCHa2R* abundance in the hindgut (Fig.3D) with trace amounts detected in other tissues.

**Figure 3.**
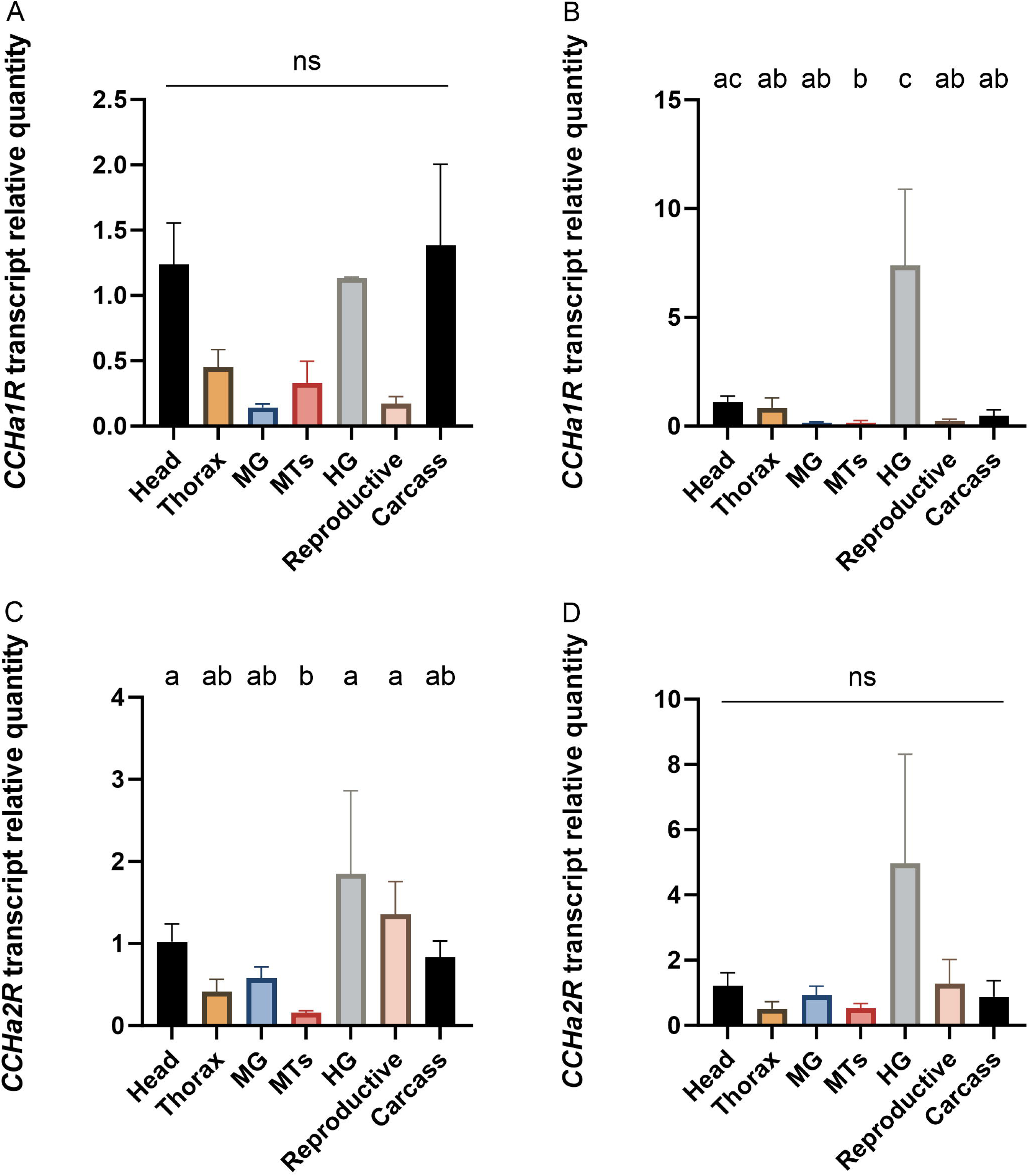
Tissue-specific expression profiles of *CCHa1R* in male (A) and female (B) along with *CCHa2R* in male (C) and female (D) 1-day-old adult *Ae. aegypti* determined by RT-qPCR. Tissue/organ abbreviations: head, thorax, midgut (MG), Malpighian tubules (MTs), hindgut (HG), reproductive system and carcass. Statistical differences are denoted with different letters, as determined by a one-way ANOVA on log-transformed data (p<0.05). Data represents the mean ± SEM (n=4).

### 3.3 Anti-*Aedae*CCHa2 antibody specificity and localization of CCHamide immunoreactivity

The binding specificity and potential cross-reaction of the affinity-purified anti-*Aedae*CCHa2 primary antibody to the neuropeptides CCHa1 and CCHa2 and other structurally distinct peptides were examined by ELISA. Results showed that, with the exception of CCHamides, other mosquito neuropeptides, including RYamide-1 and CAPA-PK1, were not recognized by the antibody. Both CCHa1 and CCHa2 were recognized similarly by the affinity purified custom antibody, despite having been generated against the latter peptide. Notably, the antibody demonstrated a slightly higher binding affinity (∼ two-fold) towards CCHa1 (EC_50_=59.29nM) compared to CCHa2 (EC_50_=134.97nM) (Supplement Figure S1).

In light of these observations regarding antibody specificity, where both CCHa1 and CCHa2 were detected similarly, whole-mount immunohistochemistry was conducted to localize CCHamide immunoreactive neurons or endocrine cells in adult mosquitoes. Approximately 15-20 immunoreactive endocrine cells were located in the posterior midgut (caudal) region adjacent to the midgut-hindgut junction known as the pyloric valve (Fig.4A). These immunoreactive cells were not observed in the negative control where antibody was preincubated overnight with CCHa2 peptide (Fig.4B). Further analysis revealed that detected immunoreactive endocrine cells in the posterior midgut likely express CCHa2 rather than CCHa1, since no immunoreactive endocrine cells were observed in the midgut following CCHa2 knockdown (Fig.4C). CCHamide immunoreactive cells were also observed in the brain and ventral nerve cord of mosquitoes. Within the brain, two pairs of CCHamide immunoreactive cells are located in the dorsal regions near the *pars intercerebralis* (PI) and *pars lateralis* (PL) (Fig.4D-E). In the terminal abdominal ganglion, three neurons were found to be CCHamide-immunoreactive (Fig.4F), while bilaterally-paired axonal projections were also observed in pre-terminal abdominal ganglia (Fig.4G). Direct tissue profiling by MALDI TOF analysis of the terminal abdominal ganglion revealed a mass match of CCHa1, whereas the ion signal responsible for CCHa2 was not detected (Fig.4H).

**Figure 4.**
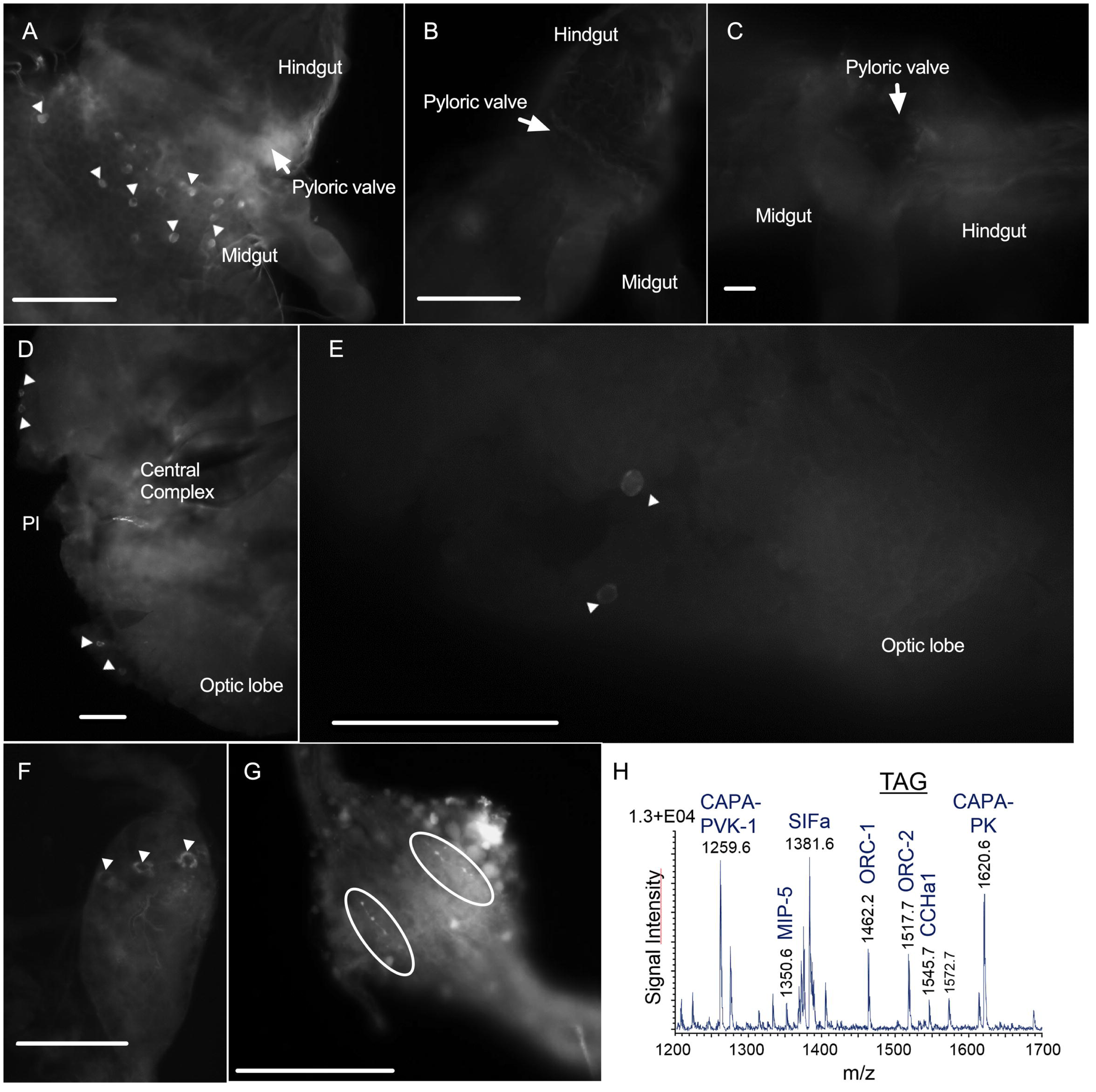
CCHamide immunoreactive cells were localized in the posterior midgut and the central nerve system of 1-day-old adult *Ae. aegypti* reared under standard laboratory conditions via custom anti-*Aedae*CCHa2 antibody. (A) Multiple immunoreactive endocrine cells (denoted by white arrowhead) localized in the posterior midgut adjacent to the pyloric valve of 1-day-old female mosquitoes. (B) Negative control midgut preparation where anti-*Aedae*CCHa2 primary antibody was pre-incubated overnight with CCHa2 (10μM). (C) CCHamide immunoreactive endocrine cells were diminished in *CCHa2* knockdown mosquitoes. (D-E) Two pairs of immunoreactive neurons (denoted by white arrowhead) localized in the *pars intercerebralis (PI)* region of 1 day old mosquitoes. (F) Immunoreactive neurons (denoted by white arrowhead) within the terminal abdominal ganglion. (G) Bilaterally-paired axonal projections (denoted by white ellipse) were detected in pre-terminal abdominal ganglia. (H) MALDI TOF mass spectrum of a preparation of the terminal abdominal ganglion which revealed a mass match of CCHa1 (m/z 1545.7). An ion signal responsible for CCHa2 was under the limit of detection and could not be detected. Similar immunolocalization was observed in both sexes. Scale bars: 50μm.

### 3.4 Mass spectrometry analysis of tissue extracts and direct tissue peptidomic profiles

Based on the immunohistochemical results, first we analyzed portions of the *pars intercerebralis*, the midgut and the terminal ganglion of the abdomen by direct tissue profiling using MALDI-TOF mass spectrometry of both sexes (Fig.5). Representative mass spectra of preparations of the *pars intercerebralis* (PI) in males and females showed an ion signal at m/z 1547.7 that is responsible for the theoretic calculated ion mass of CCHa1. A clear mass match with CCHa2 (m/z) could not be confirmed due to the ion mass similarity with myosuppressin at m/z 1247.7. Resulting direct tissue mass spectra from male and female midgut preparations are shown in Fig.5 B, D. In both male and female samples, the most abundant ion signal corresponded to CCHa1. An ion signal responsible for CCHa2 could not be detected.

**Figure 5.**
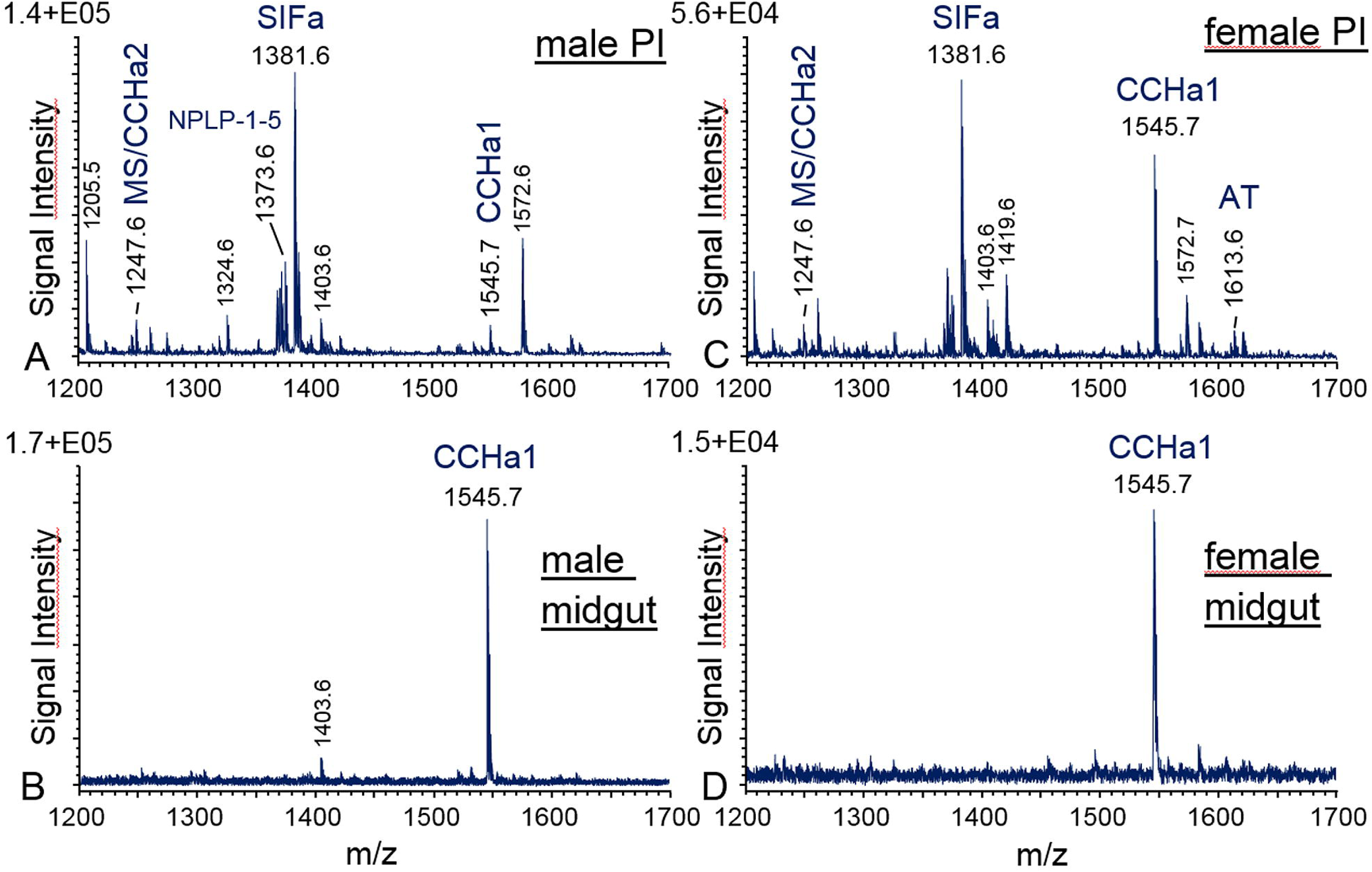
Representative MALDI-TOF mass spectra obtained from direct tissue profiling preparations of adult *Ae. aegypti* (A) male *pars intercerebralis* (PI) brain region, (B) male midgut, (C) female PI, and (D) female midgut in a mass range at m/z 1200–1700. Ion signals are marked and represent single charged peptides [M+H] ^+^. An ion signal responsible to CCHa1 was detectable in all samples. Mass spectra obtained from the PI (A, C) showed mass matches with the theoretical ion mass of CCHa2 (m/z 1247.6) which is similar to the mass of myosuppressin (MS). Additionally, the concentration of CCHa2 in midgut samples prepared by direct tissue profiling were under the limit of detection.

In the next step, we used brain and midgut extract of both sexes by Q Exactive Orbitrap MS for chemical identification of CCHamides followed by the analysis of generated fragmentation spectra using PEAKS 10.5 software package. This resulted in CCHa1 and CCHa2 sequence identification in the brain as well as in the midgut of both female (Fig.6) and male (Supplement Figure S3) mosquitoes.

**Figure 6.**
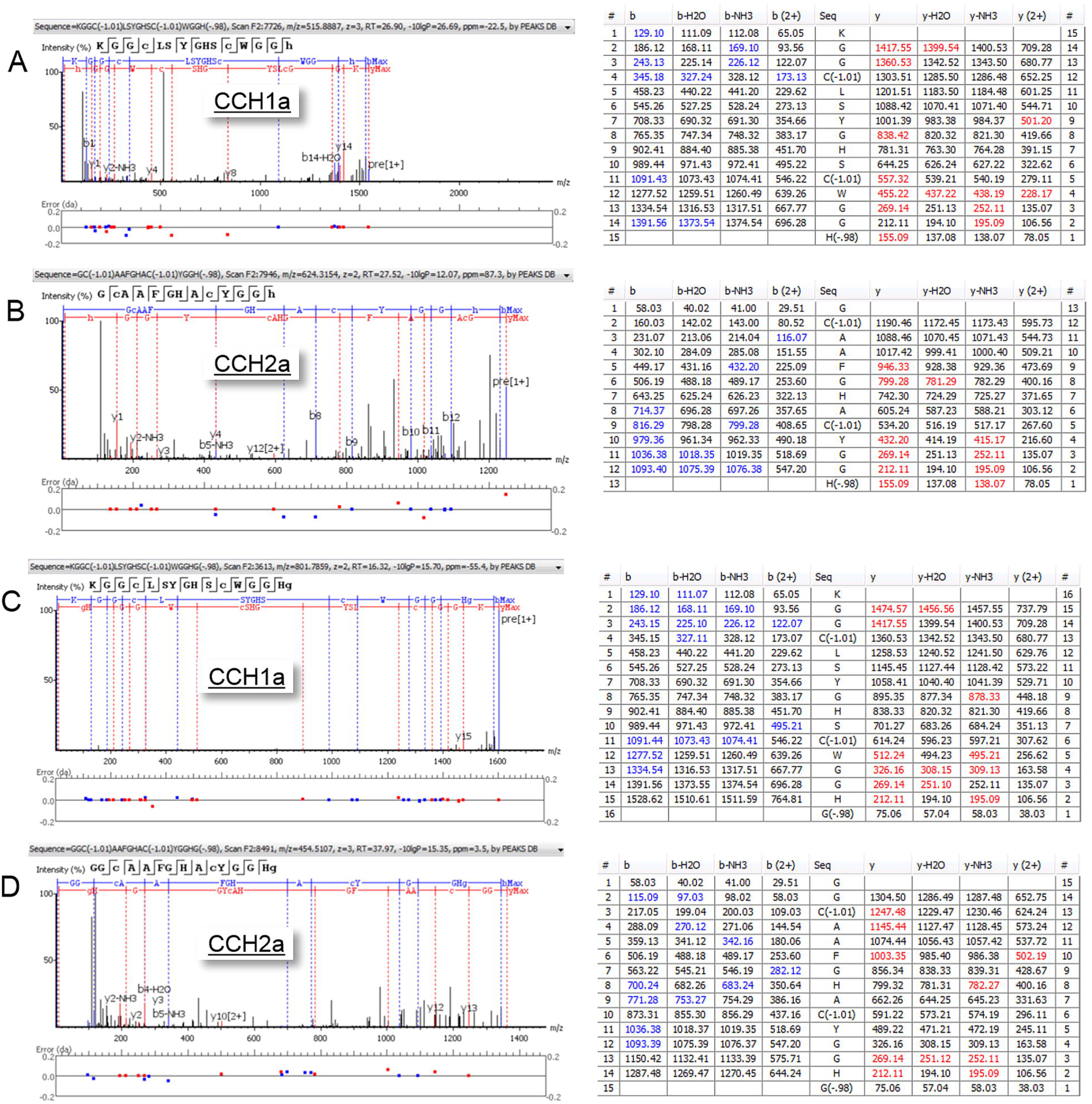
Quadrupole orbitrap MS^2^ spectra generated by ESI-Q Exactive Orbitrap MS of brain and midgut extracts of *Ae. aegypti* females for sequence identification of CCHamides. Resulting fragmentation spectra revealed the presence of CCHa1 and CCHa2 in the brain (A, B) and in midgut extracts (C, D).

### 3.5 Deorphanization of CCHa1 and CCHa2 receptors

Validation of synthesized mammalian expression constructs for each CCHamide receptor and determination of the one with the highest activation efficiency was achieved by examining luminescent responses in a heterologous expression system. Initial experiments were carried out with both CCHamide receptors to establish if heterologous receptor activity required the co-expression of the promiscuous G protein (Gα15), which indiscriminately couples a wide variety of GPCRs to calcium signaling (Offermanns and Simon, 1995). Based on these experiments, CCHa1R was co-expressed with the promiscuous G protein by using the pBudCE4.1 dual promoter vector as described previously (Wahedi et al., 2019) whereas the CCHa2R was heterologously expressed using a standard single promoter mammalian expression vector (pcDNA3.1^+^) without co-expressing the promiscuous G protein. These constructs were selected to complete further experiments including dose-response profiles because of their higher activation efficiency and greater signal-to-background noise ratio (Supplement Figure S2). Together, these data support the notion that CCHa1R may not naturally couple via calcium signaling *in vivo* whereas CCHa2R does appear to naturally utilize calcium in downstream signaling responses following ligand activation.

To deorphanize CCHamide receptors in *Ae. aegypti*, a variety of insect neuropeptides that represent diverse neuropeptide families participating in distinct physiological functions were tested to examine the sensitivity and specificity of CCHa1R (Fig.7A) and CCHa2R (Fig.7B). *Aedae*CCHa1 and *Aedae*CCHa2 were the only two neuropeptides that effectively elicited significantly elevated luminescent responses with both CCHamide receptors compared to the negative control, BSA assay media alone. Unlike the activity of CCHamides, the CCHa1R and CCHa2R were unresponsive to any other tested neuropeptides at the same saturating concentration (10^-6^M) yielding only background luminescent responses (Fig.7A-B). Both CCHamide neuropeptides elicited similar high intensity activation of CCHa2R; however, a significantly stronger response suggesting a higher binding affinity was recorded when *Aedae*CCHa1 was applied to CCHa1R relative to *Aedae*CCHa2 activity.

**Figure 7.**
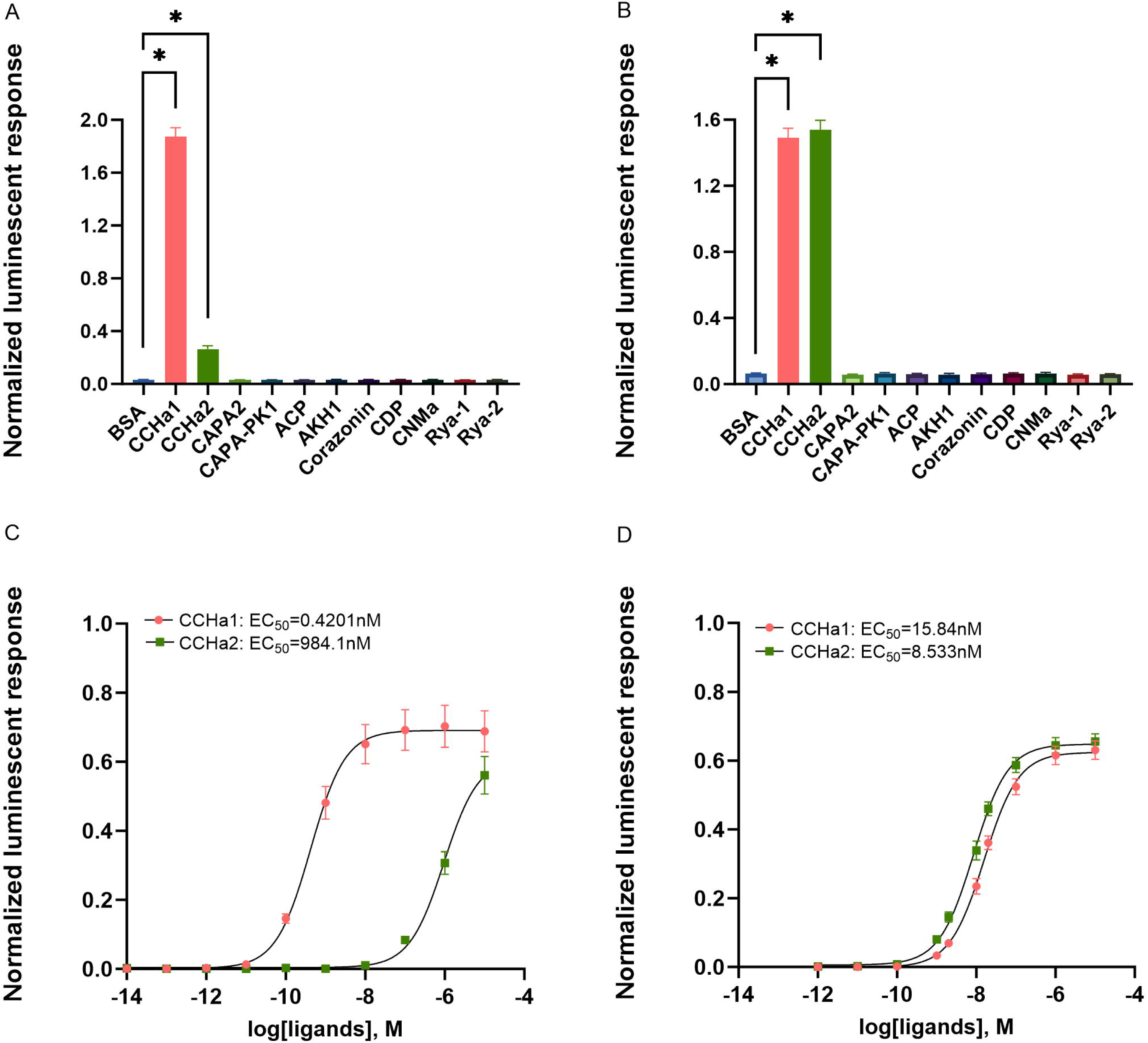
Luminescent response of CHO-K1 cells expressing *Ae. aegypti* CCHa1R and CCHa2R. CCHa1 and CCHa2 elicited a significant luminescent response activating CCHa1R (A) and CCHa2R (B). Statistical significances are denoted by an asterisk (*), as determined by a one-way ANOVA and Dunnett’s multiple comparison test (p<0.05). Dose-response curves were fitted following a sigmoidal dose-response model where CCHa1 and CCHa2 were able to activate CCHa1R (C) and CCHa2R (D) but with different binding affinities. EC_50_ values shown determined based on averages of all biological replicates (n=3).

A dose-response analysis of CCHa1R indicated a nearly 1000-fold difference in terms of receptor binding and activation efficiency between *Aedae*CCHa1 and *Aedae*CCHa2 with half maximal effective concentrations (EC_50_) of 0.4201nM and 984.1nM, respectively (Fig. 7C). The maximum luminescence responses of CCHa1R activation after application of CCHa1 was ∼ 70% relative to the response generated by ATP (the positive control), which activates endogenously expressed purinoceptors (Lajevardi and Paluzzi, 2020). Comparatively, a similar maximum luminescence response of CCHa1R was not reached in response to CCHa2 even at the highest concentration tested (10^-5^M). Next, a dose-response analysis of CCHa2R revealed a luminescence signal (representative of Ca^2+^ signaling) by both *Aedae*CCHa1 and *Aedae*CCHa2, with EC_50_ values of 15.84nM and 8.533nM, respectively (Fig. 7D) indicating CCHa2 was ∼2-fold more potent than CCHa1 in activating CCHa2R. The maximum luminescence responses of CCHa2R activation following both CCHa1 and CCHa2 activation were ∼ 60% relative to the response generated by ATP. Thus, unlike the highly selective CCHa1R that was almost 1000-fold more sensitive to *Aedae*CCHa1, CCHa2R is a promiscuous receptor that responds similarly to both endogenous CCHamides.

### 3.6 Post-blood meal CCHamide expression

Acquiring a blood meal is essential for egg production and maturation in female mosquitoes, but it also acts as a stressor challenging water and ion homeostasis. The potential influence of a blood meal on transcript abundance of CCHamides in 5 to 6-day-old female mosquitoes was examined. Results revealed that *CCHa1* and *CCHa2* transcript abundance responded differently over the 24h post-blood feeding period (Fig.8). Compared to sugar-fed control females, *CCHa*1 abundance was significantly reduced 1h after blood feeding and then recovered to the normal levels at 6h post-blood meal (PBM) and maintained this level thereafter (Fig.8A). Comparatively, *CCHa2* transcript abundance was similar to sugar-fed controls at 1h PBM and then significantly increased at 6h PBM. At 12h PBM, *CCHa2* abundance returned to the control level and did not change over the subsequent time points examined in the experiment (Fig.8B).

**Figure 8.**
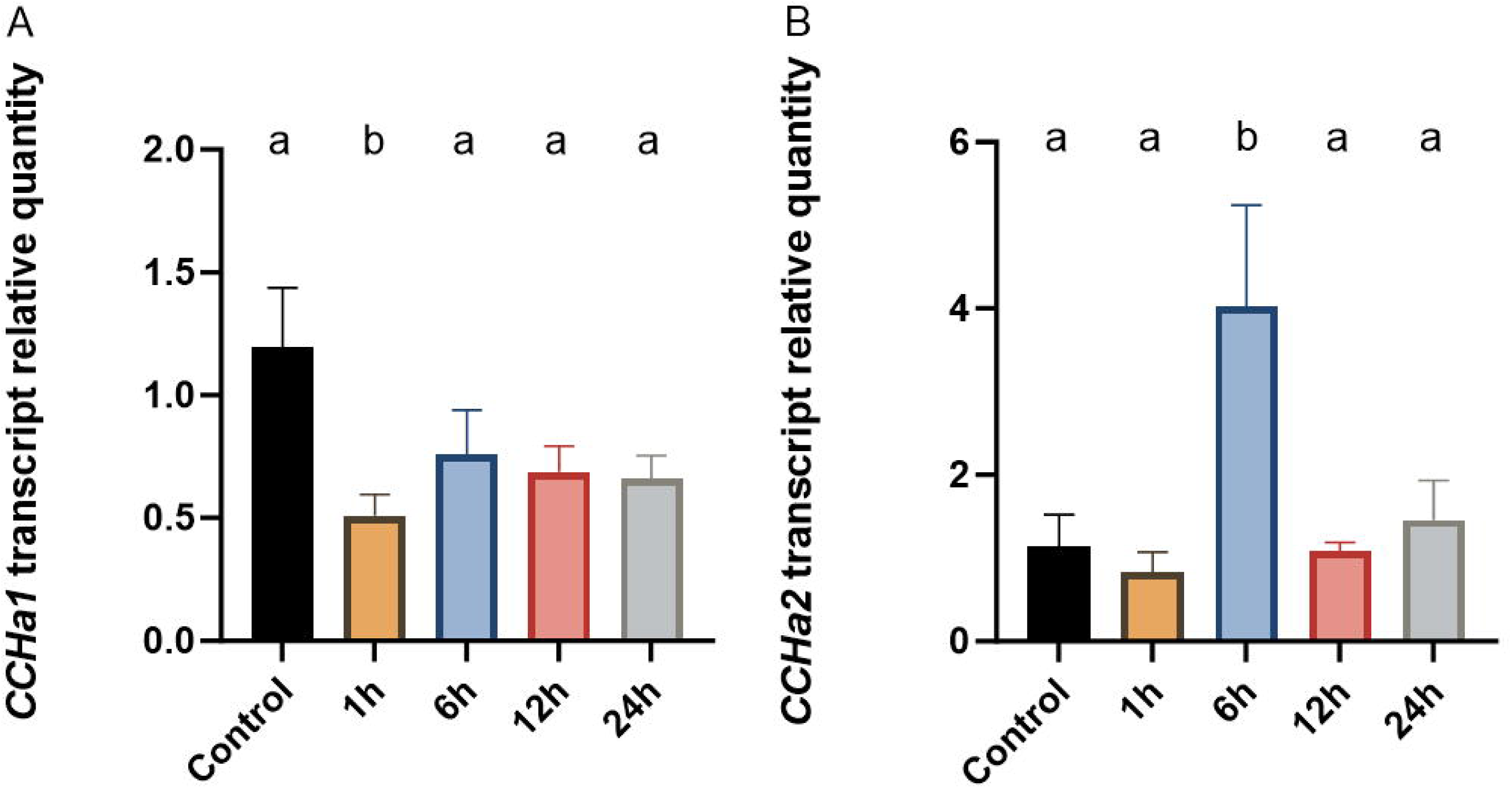
The effect of blood-feeding on the expression of *CCHa1* (A) and *CCHa2* (B) in female *Ae. aegypti*. Transcript abundance responded differently over the 24h post-blood feeding period with *CCHa1* significantly reduced at 1h PBM whereas *CCHa2* significantly increased 6h PBM. Different letters denote statistical differences, as determined by a one-way ANOVA with Dunnett’s multiple comparisons test on log-transformed data (p<0.05). Data represents the mean ± SEM (n=5).

### 3.7 CCHamide transcript expression during starvation stress

Based on studies in other insects that indicated that the function of CCHa2 and CCHa2R was tightly associated with food consumption and feeding behaviours (Ren et al., 2015; Sano et al., 2015; Shahid et al., 2021; Titos and Rogulja, 2020; Zhu et al., 2022), the influence of starvation stress on the abundance of *CCHa2* and *CCHa2R* in 1-day-old male and female mosquitoes was examined. The 24h duration was selected since preliminary experiments indicated that a longer duration (48 hours) of starvation appeared to exert extremely high stress, resulting in the death of the vast majority of experimental mosquitoes. Following this 24h stress experimental regime, neither starvation alone or starvation combined with water deprivation impacted the transcript abundance of *CCHa2* and *CCHa2R* (Fig.9A-D). Immunohistochemistry was also conducted on 24h starved and desiccated mosquitoes where a similar number of CCHamide immunoreactive endocrine cells were found in the caudal region of the posterior midgut as observed in mosquitoes under non-stressed conditions (Fig.9E-F). Further, while maintaining image acquisition parameters constant, the intensity of CCHamide immunoreactive staining of posterior midgut endocrine cells was similar between sugar-fed control and starvation– and water-stressed experimental groups.

**Figure 9.**
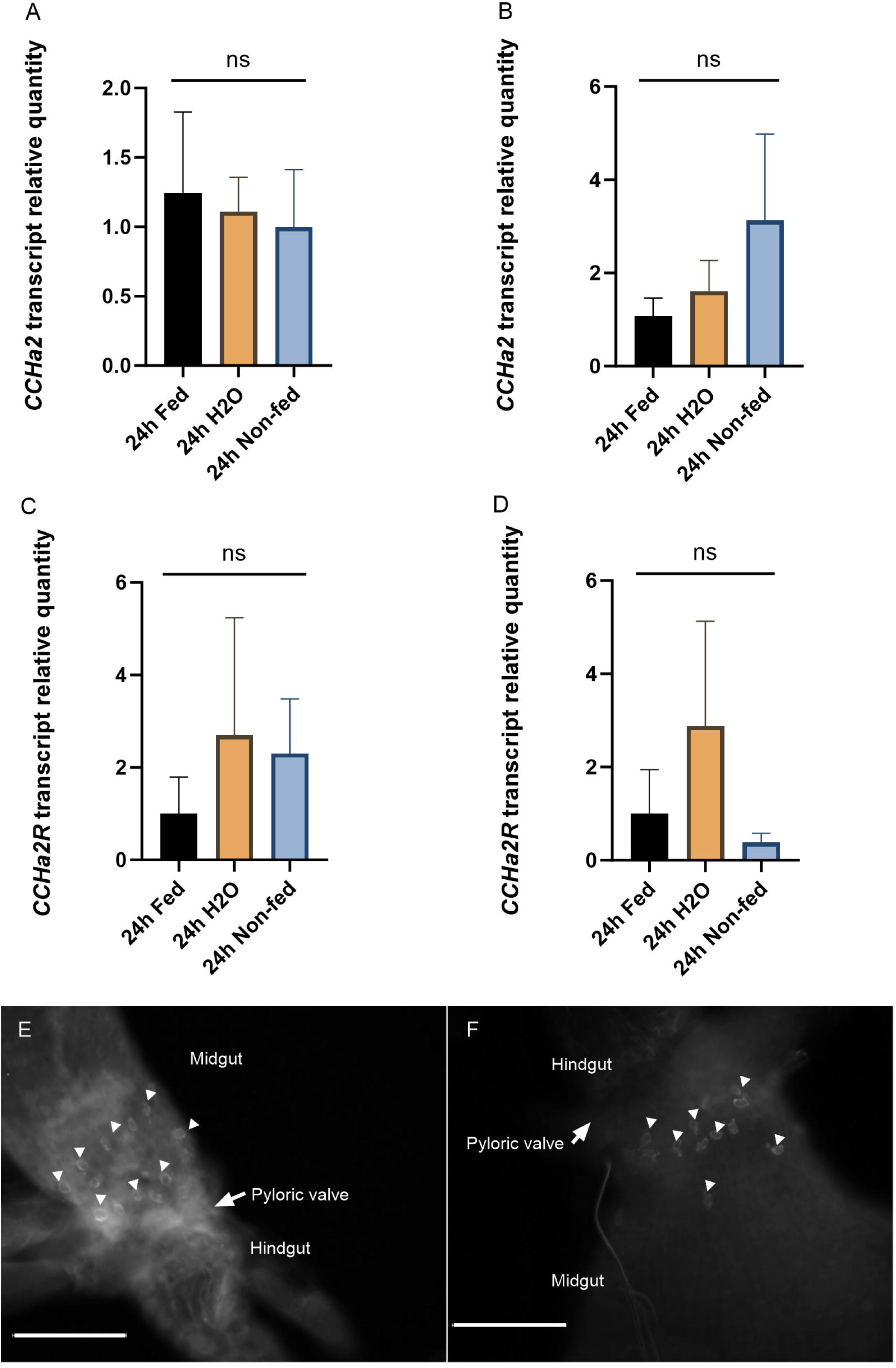
Single day (24h) water deprivation and/or starvation does not influence expression of *CCHa2* in male (A) and female (B) mosquitoes, and *CCHa2R* in male (C) and female (D) mosquitoes or lead to changes in CCHamide immunoreactivity in midgut enteroendocrine cells (E-F). In A-D, statistical analysis involved a one-way ANOVA with Tukey’s multiple comparisons test on log-transformed data (p<0.05). Data represents the mean ± SEM (n=3). CCHamide immunoreactive endocrine cells in the posterior midgut of *Ae. aegypti* raised under standard laboratory conditions where animals were sucrose fed *ad libitum* (E) and after 24h combined starvation and desiccation (F). Multiple immunoreactive endocrine cells (denoted by white arrowhead) with similar number and intensity of CCHamide immunoreactivity as observed in sucrose-fed mosquitoes. Scale bars: 50μm.

**Figure 10.**
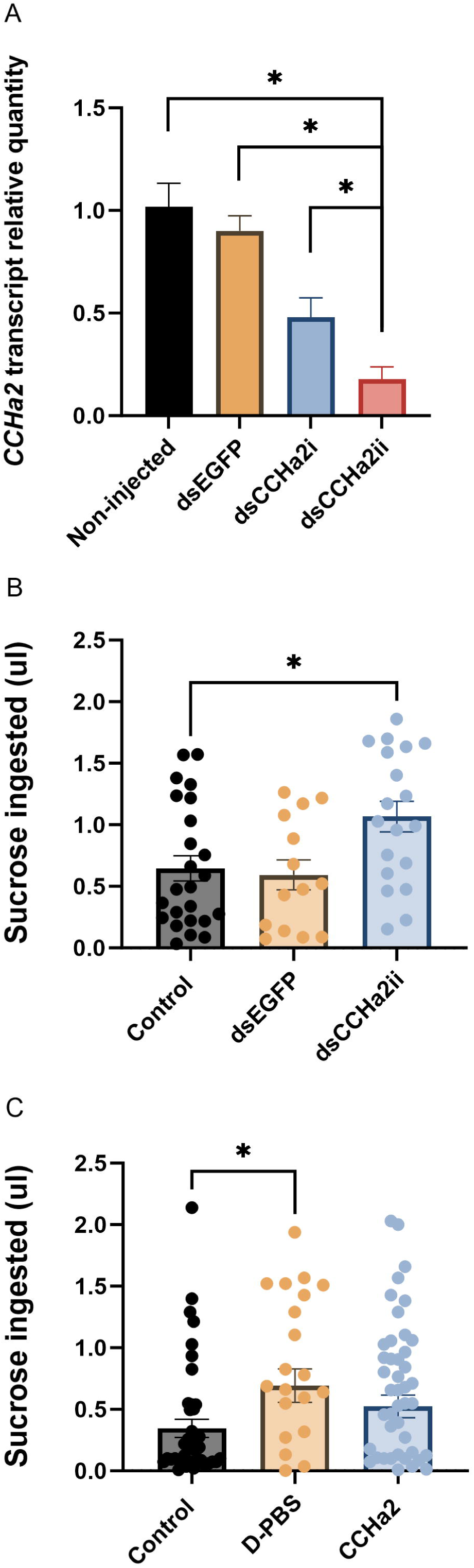
*CCHa2* knockdown by RNAi increases sucrose ingested over 24h by female adult *A. aegypti* mosquitoes. (A) One-day-old female mosquitoes were injected with synthesized dsRNAs targeting either EGFP (ds*EGFP*), 5’ UTR (dsCCHa2i) or 3’ UTR (ds*CCHa2ii*) of *A. aegypti* CCHa2 gene. Knockdown efficiency was assessed by RT-qPCR analysis on 4-day-old female mosquitoes (3 days post dsRNA injection). Statistical differences are denoted with an asterisk (*), as determined by a one-way ANOVA with Tukey’s multiple comparisons test on log-transformed data (p<0.05). Data represent the mean ± SEM (n=3). (B) Using the CCHa2 dsRNA that led to the greatest knockdown efficiency (dsCCHa2ii), sucrose feeding by adult female mosquitoes was measured over a 24h period. Knockdown of CCHa2 led to a significant increase (∼two-fold) of ingested sucrose by adult females. (C) Injection of female mosquitoes with synthetic CCHa2 did not influence the volume of sucrose solution ingested over a 24h period, but it did not undergo the same increase as observed in the saline (D-PBS) injected mosquitoes. Statistical differences are denoted with an asterisk (*), as determined by a one-way ANOVA with Dunnett’s multiple comparisons test (B-C) (p<0.05).

### 3.8 Verification of efficient CCHa2 knockdown by RNAi

RNAi targeting *CCHa2* in adult *A. aegypti* was achieved by direct injection of synthesized dsCCHa2. In comparison to the non-injected control group, there was no change in *CCHa2* transcript abundance in the ds*EGFP*-injected group. Injection of ds*CCHa2* targeting either region of the *CCHa2* gene led to a significant reduction in *CCHa2* transcript expression compared to the control females (Fig.10A). Moreover, the knockdown efficiency was higher for the ds*CCHa2* targeting the 3’ UTR (ds*CCHa2ii*), where *CCHa2* transcript abundance was reduced by 90%, compared to the dsCCHa2 (ds*CCHa2i*) spanning the 5’ UTR, where *CCHa2* transcript was knocked down by ∼50%. Due to its superior knockdown performance, ds*CCHa2* targeting 3’UTR of the *CCHa2* gene was used for subsequent immunohistochemistry and CAFE assay.

### 3.9 Effects of CCHa2 signaling in sucrose feeding

To examine whether CCHa2 is involved in regulating the sugar-feeding behaviour of adult female *A. aegypti*, a capillary feeder (café) assay was conducted to measure sucrose intake in the control, *CCHa2* knockdown and CCHa2-injected group. The average sucrose ingested by control (non-injected) female mosquitoes over a 24h period was similar to animals injected with ds*EGFP* (Fig.10B). However, relative to these two control groups, decreasing the abundance of *CCHa2* transcript in female mosquitoes by ds*CCHa2* injection resulted in a significant increase in food intake over a 24h period, which was nearly two-fold higher compared to the control mosquitoes. In comparison to the knockdown group, elevating the CCHa2 neuropeptide in the haemolymph of female mosquitoes did not alter the feeding amount since no significant difference in sucrose intake was observed between the control and *Aedae*CCHa2 injected group (Fig.10C). However, in comparison to the saline (D-PBS) injected mosquitoes, which resulted in significantly increased feeding, no such increase in sucrose intake was observed in the *Aedae*CCHa2 injected group, suggesting this peptide might inhibit sucrose intake.

## 4. Discussion

CCHamide neuropeptide sequences were noted over two decades ago (Zdárek et al., 2000), and biochemically isolated and sequenced a decade later where they were identified as endogenous ligands for orphan receptors, CCHa1R and CCHa2R in *Drosophila* (Ida et al., 2012). However, beyond studies in the fruit fly *D. melanogaster* and a few other insects, research on CCHamides is limited. This has kept CCHamides and their associated receptors uncharacterized among many other insects, including the mosquito, *Ae. aegypti*. In this study, we have functionally deorphanized two CCHamide receptors, examined both developmental and spatial expression profiles of transcripts encoding CCHamides and their receptors, localized the distribution of CCHamide immunoreactivity in adults, and used RNAi to knockdown *CCHa2* expression in order to elucidate potential physiological roles in the mosquito, *Ae. aegypti*.

### 4.1 Developmental expression of CCHamide and CCHamide receptor transcripts

CCHamide transcripts were found to be enriched in mosquito adults with *CCHa2* levels being more prominent in earlier adulthood while *CCHa1* was enriched in older adults. Additionally, the *CCHa1R* transcript abundance also showed enrichment during adult stages, while the highest abundance of *CCHa2R* was found in the late pupal stage. This pattern is generally consistent with findings in *D. melanogaster*, where qPCR results demonstrated *CCHa1R* was enriched in male adults, while pupae expressed the most *CCHa2R* transcript along with adult male flies (Li et al., 2013). Notably, nutrient-dependent factors can lead to sex-specific differences since *CCHa2* decreased in *D. melanogaster* male larvae raised on a protein-rich diet while no change was observed in female larvae (Millington et al., 2021). Relatedly, a sex-specific function of CCHamides in adult mosquitoes may be possible since both peptide and receptor transcripts were generally more abundant in the male mosquitoes, apart from *CCHa2R*. Despite *Ae. aegypti* and *D. melanogaster* diverged around 260 million years ago (Arensburger et al., 2010), the regulatory role of CCHa2 uncovered in fly development provides insight for investigating its potential role in mosquitoes given the similar expression patterns (Strand et al., 2016; Wiegmann et al., 2003). Future studies could investigate if CCHa2 is involved in larval growth and pupariation, and whether it facilitates these events directly or indirectly through other neuropeptides or signaling pathways, for example, ILPs (Ren et al., 2015; Sano et al., 2015).

*CCHa2* had increased abundance in the late pupa stage just before eclosion, which suggests CCHa2 may play a role in this process or events in preparation for eclosion. In addition, its corresponding receptor (*CCHa2R*) had the highest abundance in the late-stage pupa, which differs from the expression profile reported for the homologous receptor in the pea aphid, where highest transcript level was found in earliest developmental stage (Shahid et al., 2021). Taken together, the findings in mosquito suggest that this receptor may be functional in the late pupae or early adult stage facilitating downstream responses through CCHa2-CCHa2R signaling.

Until now, there is no evidence that supports CCHa1-CCHa1R signaling is involved in insect growth, development and eclosion, but rather, it was linked to specific diet and associated with arousability and olfactory responses under starvation stress in flies (Farhan et al., 2013; Titos and Rogulja, 2020). Considering the high enrichment of *CCHa1* in older adult-stage mosquitoes, this may suggest that CCHa1 holds a more prominent function in the adult stage rather than in development as demonstrated for CCHa2-CCHa2R signaling.

### 4.2 Midgut is a major source for CCHamide production

Tissue expression profiles revealed that the mosquito midgut is a primary source producing CCHa1 and CCHa2 peptides alongside the CNS. CCHamide immunoreactivity was evident as multiple enteroendocrine cells localized in the posterior (caudal) region of the posterior midgut adjacent to the pyloric valve of both male and female adult mosquitoes. In support of this CCHa immunoreactivity, direct tissue profiling and tissue extract analysis by mass spectrometrical methods validated the midgut as a bona fide source of both CCHa1 and CCHa2 in the adult *Ae. aegypti* mosquito. Insect midgut is an important organ responsible for food digestion, nutrient absorption and water and ion homeostasis (Caccia et al., 2019; Cui and Franz, 2020). Multiple neuropeptides with roles in various physiological processes are known to be localized within intestinal endocrine cells in the insect midgut (Wu et al., 2020). Studies have revealed that at least 10 kinds of neuropeptides are observed in the midgut of *Drosophila* (Chen et al., 2016; Reiher et al., 2011) and these gut-derived neuropeptides are involved in various physiological processes including development and ecdysis, feeding and nutritional status-related functions, nutrient and ion transport as well as intestinal muscle contraction (Abou El Asrar et al., 2020; Gelman et al., 1991; Wu et al., 2020).

The expression profile of CCHamide transcripts in female *A. aegypti* was dynamic since *CCHa1* abundance was significantly reduced at 1h PBM, while *CCHa2* increased significantly at 6h PBM. Since 1h PBM spans the period where more active excretory processes occur to remove excessive ions and water, and *CCHa1R* was enriched in the hindgut that is tightly associated with these processes, CCHa1-CCHa1R signaling may participate in maintaining water-ion homeostasis. As gut-derived neuropeptides, CCHamides were earlier found to be associated with feeding and feeding-related activity, as *CCHa1* expression increased following ingestion of protein-rich food, but not in response to other nutritional substances like sugars or fats (Titos and Rogulja, 2020). Interestingly, besides female mosquitoes consuming a protein-rich blood meal for egg maturation during each gonotrophic cycle, both female and male mosquitoes feed on plant nectar (Barredo and DeGennaro, 2020). Considering this, what function (if any) does CCHa1 possess in mosquitoes of both sexes in relation to nectar-feeding and might *CCHa1* be modulated through factors other than protein ingestion? In *D. melanogaster*, it was revealed that the abundance of *CCHa1R* is influenced by starvation stress in flies (Farhan et al., 2013), suggesting CCHa1-CCHa1R signaling may regulate foraging behaviour of mosquitoes.

In addition to the factors influencing expression of CCHamides and their receptors discussed above, *CCHa2* abundance can be altered by nutritional conditions since starved fruit fly larvae had significantly lower levels of *CCHa2* transcript (Sano et al., 2015) and male larvae also had significantly lower levels when reared on a protein-rich diet (Millington et al., 2021). However, results in the current study on *A. aegypti* did not align with these earlier findings since the abundance of *CCHa2* and *CCHa2R* was not affected by starvation/desiccation stresses. In support of this conclusion, a similar number of CCHamide immunoreactive endocrine cells having comparable staining intensities were localized within the same posterior (caudal) region of the posterior midgut in both sucrose-fed and starved/desiccated mosquitoes. Previous studies in *Drosophila* indicated that CCHa2 is a sugar-responsive neuropeptide since *CCHa2* knockout leads to reduced food-seeking time and food intake (Ren et al., 2015). In the current study, CCHa2 was also demonstrated as a sugar-responsive neuropeptide in *A. aegypti*, but surprisingly, it appeared to have an opposite regulatory function. Knocking down *CCHa2* transcript abundance in female adult mosquitoes resulted in a significantly higher sucrose intake. Increasing CCHa2 neuropeptide in the haemolymph (by injection) had no effect on sucrose ingestion (i.e., did not decrease sucrose intake compared to the control group), which may be due to endogenous CCHa2 saturating CCHa2R available. However, saline injected mosquitoes exhibited significantly greater sucrose intake under the same condition, and this may explain to some extent that the elevated CCHa2 neuropeptide in the haemolymph suppresses the food intake of female mosquitoes, which align with the complementary results obtained from *CCHa2* knockdown mosquitoes where sucrose feeding increased. Therefore, these results support the notion that CCHa2 acts as a sugar-feeding repressor in *A. aegypti*.

### 4.3 The central nervous system is an additional source for CCHamide neuropeptides

In addition to the high expression of CCHamide transcripts in the midgut and their detection by mass spectrometric and immunohistochemical methods, CCHamide immunoreactive neurons and processes were localized within the central nervous system, including the brain and ventral nerve cord of mosquitoes. In the cricket, *G. bimaculatus*, *CCHa2* transcript was similarly detected in numerous tissues including the midgut as well as the brain, thoracic and abdominal ganglia (Zhu et al., 2022). Diverse biological processes in insects are regulated by factors derived from the ventral nerve cord, such as development, water and ion homeostasis, hindgut transepithelial transport and motility, mating and oviposition (Nässel, 1996; Raabe, 2012; Weevers, 1985). Knowledge on neuropeptides produced and released in similar regions of the abdominal ganglia could shed light on the role of CCHamides in mosquitoes. Neuropeptides expressed in the terminal abdominal ganglia (TAG) are known to mediate gut muscle contraction that may be associated with nutrient digestion, uptake of water, ion reabsorption, and mediating reproduction (Nässel, 1996, 2002). For example, immunoreactivity of FMRFamide-related peptides (FaRPs) was observed in the posterior region of the terminal abdominal ganglion (TAG) in the migratory locust, which project into the ventral ovipositor nerve and the receptaculum seminis nerve. The authors deduced that FaRPs affect the amplitude of neurally evoked spermathecal contractions (Clark and Lange, 2002). Moreover, six proctolin immunoreactive neurons were observed in the anterior region of the TAG in grasshoppers and proctolin is also known to enhance muscle contraction of the midgut and hindgut of *R. prolixus* (Keshishian and O’Shea, 1985; Orchard et al., 2011). Since CCHamide immunoreactive neurons have been detected in the mosquito TAG with axonal processes projected posteriorly towards the hindgut and reproductive organs, could CCHamides intervene in these or related physiological processes? This and other outstanding questions will require additional investigation of the CCHamide signaling systems.

### 4.4 Deorphanization of *Ae. aegypti* CCHamide receptors

One of the main objectives of this study was achieved since both *Aedae*CCHamide receptors were deorphanized with *Aedae*CCHa1 and *Aedae*CCHa2 confirmed as the most potent ligands specifically activating *Aedae*CCHa1R and *Aedae*CCHa2R, respectively. Unlike the other peptidergic ligands tested that yielded no response, CCHa1 and CCHa2 exhibited high activity on their receptors and were able to trigger downstream effects in the form of Ca^2+^ release. One key difference observed was that CCHa1R had significantly higher sensitivity to CCHa1 relative to CCHa2, while both neuropeptides were similarly effective in activating CCHa2R. This indicates that the *Ae. aegypti* CCHa1R is quite selective for its native ligand, CCHa1 whereas the CCHa2R is rather promiscuous and does not differentiate between the two CCHamide peptides in mosquitoes. It was earlier reported that both the disulfide bond and the amide group at the C-terminus of *Drosophila* CCHa1 and CCHa2 are essential for activating their corresponding receptors (Ida et al., 2012). While the core tridecapeptide consensus CCHamide sequence of *D. melanogaster* is XCXXYGHXCXGXHamide, in the mosquito *Ae. aegypti* the consensus is GCXXXGHXCXGGHamide with greater identity including a conserved C-terminal tripeptide motif, that could explain why *Aedae*CCHa1 and *Aedae*CCHa2 can activate both receptors including the highly promiscuous CCHa2R. Comparatively, our findings revealed that CCHa1R was much more selective in terms of differentiating between these closely related neuropeptides as CCHa1 exhibited a roughly 2000-fold higher activation efficiency than CCHa2. Although CCHa2R was unable to distinguish between high doses of CCHa1 and CCHa2, the dose-response analysis revealed that CCHa2 was only ∼3-fold more potent than CCHa1. While this heterologous data should be considered carefully, it raises the notion that, besides CCHa2 mediated activity, CCHa1 may under certain conditions activate CCHa2R in the mosquito *Ae. aegypti*. Further examining this potential crosstalk between CCHamide signaling could involve similar studies using *Culex* and *Anopheles* mosquito species since members of this group contain CCHamide neuropeptides resembling the *D. melanogaster* sequence where the penultimate residue of CCHa1 is an alanine as opposed to a glycine as seen in *Ae. aegypti* (Hansen et al., 2011). On the other hand, CCHa2 is unlikely to compete with CCHa1 as a bonafide ligand of the CCHa1R *in vivo* since CCHa1 was over three orders of magnitude more potent on the CCHa1R. At the moment, this fact CCHa2R responds similarly to both endogenous CCHamides is unusual considering that the two receptors in *D. melanogaster* are specifically activated by either CCHamide-1 (CG30106), or CCHamide-2 (CG14593) with each receptor exhibiting at least 2-3 orders of magnitude selectivity with their respective peptide (Hansen et al., 2011; Ida et al., 2012). Thus, while in *D. melanogaster* there are two distinct CCHamide signaling systems, such a conclusion cannot be reached for the CCHamides in the mosquito *A. aegypti*.

### 4.5 Concluding remarks and future directions

This study reported the developmental and spatial expression profiles of *CCHa1*, *CCHa2*, *CCHa1R* and *CCHa2R* in the mosquito *Ae. aegypti*. Further, peptidomics was used to validate the amino acid sequences of mature CCHamide neuropeptides by mass spectrometry along with their organ-specific distribution. Relatedly, immunohistochemistry localized CCHamide immunoreactive endocrine cells and neurons, and a heterologous assay was used to deorphanize the two CCHamide receptors, CCHa1R and CCHa2R. Our results support that CCHa2-CCHa2R signaling is involved in regulating feeding behaviour of mosquitoes since CCHa2 acts as a sugar-feeding repressor under non-stressed conditions. However, details are unclear regarding any potential association between CCHamides and blood-meal feeding by female mosquitoes, thus further research is needed to clarify whether CCHamides influence mosquito reproduction or development. One means of answering these questions could be using reverse genetic tools such as CRISPR-Cas9 to knockout CCHamides or their receptors and examine any resultant abnormal phenotypes of these mutant mosquitoes. Nonetheless, the current study has advanced our understanding of mosquito neuropeptide signaling and the physiological roles that are governed by CCHamides. As mosquitoes transmit diseases related to their blood-feeding behaviour, studying neuropeptides involved in regulating foraging and feeding processes might provide insight for new strategies or specific biorational compounds useful in pest control.

## Acknowledgments

This work was supported by a Natural Sciences and Engineering Research Council of Canada (NSERC) Discovery Grant and an Ontario Ministry of Research Innovation Early Researcher Award to J. P. Paluzzi.

## CRediT authorship contribution statement

**Jinghan Tan:** Conceptualization, Data curation, Formal analysis, Investigation, Methodology, Resources, Software, Validation, Visualization, Writing – original draft, Writing – review & editing. **Susanne Neupert:** Investigation, Validation, Writing – review & editing. **Jean-Paul Paluzzi:** Conceptualization, Resources, Funding acquisition, Project administration, Supervision, Writing – review & editing.

## Data availability

Data will be made available on request.

