## Supplementary information for "Functional characterization of CCHamides and deorphanization of their receptors in the yellow fever mosquito, *Aedes aegypti*"

**Table S1.** Information for oligonucleotides used for preparing CCHamide receptor construct used in heterologous functional assay and dsRNA synthesis.

| Oligo Name | Sequences (5’ > 3’) | Function |
|---|---|---|
| CCHa1R-Res-756F | AAAAAGCTTgccaccATGTTGGACCCAGGG | ORF cloning of CCHa1R for functional assay |
| CCHa1R-Res-2154R | AAATCTAGAATGTGAAAATAGTGCTCCTA | ORF cloning of CCHa1R for functional assay |
| CCHa2R-Res-29F | AAGCTTgccaccATGGAACTGTGGCGCTTGATAC | ORF cloning of CCHa2R for functional assay |
| CCHa2R-Res-1336R | TCTAGAGAAATACACCATCAGCCGCAC | ORF cloning of CCHa2R for functional assay |
| CCHa2 RNAi-273F | GTGGAGTGGAAGATAGTGACCG | dsRNA synthesis targeting AedaeCCHa2 (Target 1) |
| CCHa2 RNAi-610R | TTCTCCGACAGTGACGATGATC | dsRNA synthesis targeting AedaeCCHa2 (Target 1) |
| CCHa2 RNAi-968F | AAGAACTCTTCCCCATTCCGAC | dsRNA synthesis targeting AedaeCCHa2 (Target 2) |
| CCHa2 RNAi-1450R | GTTTTGCCGGACTGTCTTCTTC | dsRNA synthesis targeting AedaeCCHa2 (Target 2) |
| EGFP-RNAi-F2 | ACTCGTGACCACCCTGACCTACG | dsRNA synthesis targeting EGFP gene |
| EGFP-RNAi-R2 | AGATCTTGAAGTTCACCTTGATGCC | dsRNA synthesis targeting EGFP gene |

**Table S2.** Information for oligonucleotides used in RT-qPCR analysis of *CCHa1*, *CCHa2*, *CCHa1R* and *CCHa2R* to determine developmental and adult tissue-specific transcript levels.

| Oligo Name | Sequences (5’ > 3’) | Function | Amplification efficiency (%) | Product size |
|---|---|---|---|---|
| CCHa1-737F | CTCCGTCTGTTTCGTTAGTGGA | RT-qPCR amplification of *Aedae*CCHa1 | 101.6% | 164bp |
| CCHa1-901R | GTTGTCGAAGCAGATCATCACG | | | |
| CCHa2-247F | AGTAAAACATTCGATGACGCCTCC | RT-qPCR amplification of *Aedae*CCHa2 | 95.2% | 227bp |
| CCHa2-474R | CAGTGATGGTGCCTGTTGC | | | |
| CCHa1R-1183F | CGTCAATTCTCTACACGGTGGA | RT-qPCR amplification of AedaeCCHa1R | 111.4% | 74bp |
| CCHa1R-1255R | TCCTTGATGAACTCGGACGAAG | | | |
| CCHa2R-334F | AACACCTACATTTTCTCGCTGG | RT-qPCR amplification of AedaeCCHa2R | 89.5% | 226bp |
| CCHa2R-560R | GTCTGCAGCTTTCGTAAAGG | | | |

**Table S3.** List and primary structure of several insect neuropeptides tested for functional activation of the mosquito CCHamide receptors using heterologous assay.

| Peptide name | Peptide sequences |
|---|---|
| *Aedae*CCHamide1 | KGGCLSYGHSCWGGH-$NH_2$ |
| *Aedae*CCHamide2 | GCAAFGHACYGGH-$NH_2$ |
| *Aedae*CAPA2 | pQGLVPFPRV-$NH_2$ |
| *Aedae*CAPA-PK1 (Pyrokinin1) | AGNSGANSGMWFGPRL-$NH_2$ |
| *Aedae*ACP | pQVTFSRDWNA-$NH_2$ |
| *Aedae*AKH1 | pQLTFTPSW-$NH_2$ |
| Corazonin | pQTFQYSRGWTN-$NH_2$ |
| CDP | NPFHSWG-$NH_2$ |
| *Aedae*CNMa | YMSLCHFKLCNM -$NH_2$ |
| *Aedae*RYamide1 | PVFFVASRY-$NH_2$ |
| *Aedae*RYamide2 | NDRFFLGSRY-$NH_2$ |

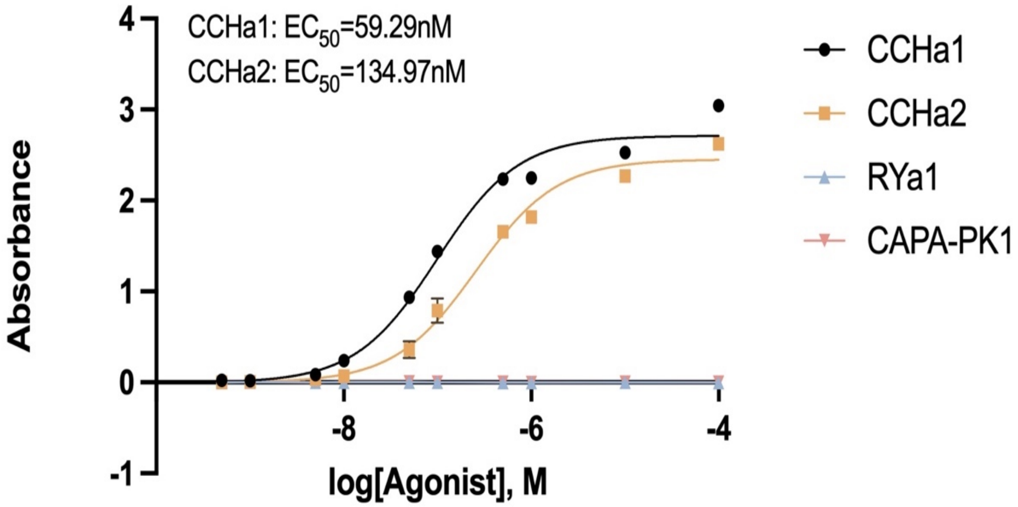


**Figure S1. Binding affinities of custom antibody to synthetic neuropeptides CCHa1, CCHa2, RYa-1 and CAPA-PK1 measured by ELISA test.** The antibody anti-*Aedae*CCHa2 was designed to target the *Aedae*CCHa2 sequence, GCQAYGHVCYGGH-$NH_2$. The antibody had high affinity to *Aedae*CCHa1 ($EC_{50}$ = 59.29nM) and *Aedae*CCHa2 ($EC_{50}$ = 134.97nM) and did not recognize other neuropeptides tested including *Aedae*RYa-1 and *Aedae*CAPA-PK1. $EC_{50}$ values represent the average values of multiple replicates, which were analyzed by sigmoidal dose-response model.

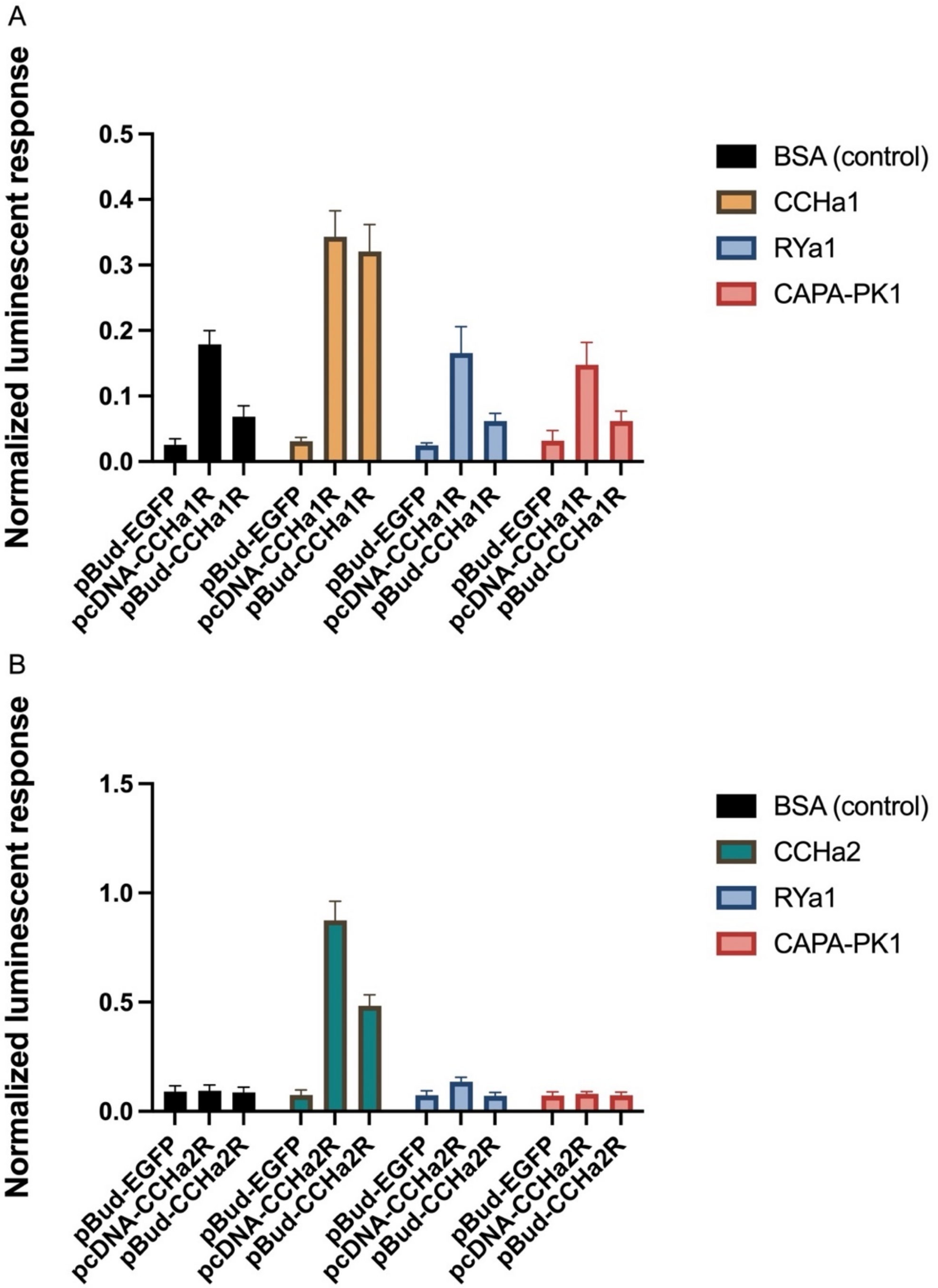


**Figure S2. Normalized luminescent response of CHO-K1 cells stably expressing aequorin and transiently expressing either EGFP, *Ae. aegypti* CCHa1R (A) or CCHa2R (B) in pcDNA3.1+ or pBudCE4.1 dual promoter vector containing the murine promiscuous Galpha15.** Assay media (BSA) alone and different ligands were applied to cells expressing different expression constructs to validate the CCHamide receptor activity by comparing the luminescent responses generated via receptor activation. The figure is taken from a single replicate but is representative of results obtained in repeated experimental trials (n=3).

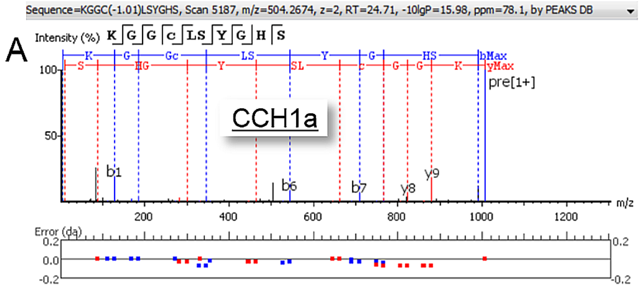


| # | b | b-H2O | b-NH3 | b (2+) | Seq | y | y-H2O | y-NH3 | y (2+) | # |
|---|---|---|---|---|---|---|---|---|---|---|
| 1 | 129.10 | 111.09 | 112.08 | 65.05 | K | | | | | 10 |
| 2 | 186.12 | 168.11 | 169.10 | 93.56 | G | 879.43 | 861.42 | 862.41 | 440.18 | 9 |
| 3 | 243.15 | 225.14 | 226.12 | 122.07 | G | 822.41 | 804.32 | 805.38 | 411.67 | 8 |
| 4 | 345.22 | 327.14 | 328.20 | 173.07 | C(-1.01) | 765.39 | 747.30 | 748.36 | 383.16 | 7 |
| 5 | 458.23 | 440.22 | 441.20 | 229.62 | L | 663.31 | 645.31 | 646.28 | 332.16 | 6 |
| 6 | 545.30 | 527.29 | 528.24 | 273.14 | S | 550.23 | 532.22 | 533.20 | 275.61 | 5 |
| 7 | 708.37 | 690.36 | 691.31 | 354.69 | Y | 463.23 | 445.22 | 446.20 | 232.10 | 4 |
| 8 | 765.39 | 747.34 | 748.36 | 383.17 | G | 300.17 | 282.16 | 283.14 | 150.57 | 3 |
| 9 | 902.41 | 884.40 | 885.38 | 451.70 | H | 243.11 | 225.10 | 226.08 | 122.05 | 2 |
| 10 | | | | | S | 106.05 | 88.04 | 89.02 | 53.52 | 1 |

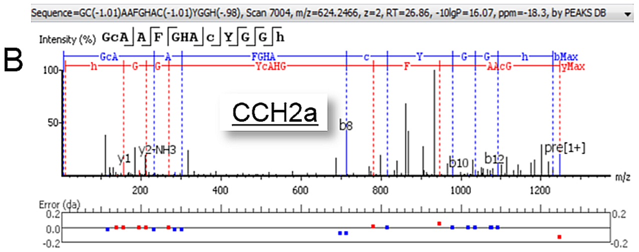


| # | b | b-H2O | b-NH3 | b (2+) | Seq | y | y-H2O | y-NH3 | y (2+) | # |
|---|---|---|---|---|---|---|---|---|---|---|
| 1 | 58.03 | 40.02 | 41.00 | 29.51 | G | | | | | 13 |
| 2 | 160.03 | 142.02 | 143.00 | 80.52 | C(-1.01) | 1190.46 | 1172.45 | 1173.43 | 595.73 | 12 |
| 3 | 231.10 | 213.06 | 214.04 | 116.07 | A | 1088.46 | 1070.45 | 1071.43 | 544.73 | 11 |
| 4 | 302.13 | 284.12 | 285.08 | 151.55 | A | 1017.42 | 999.41 | 1000.40 | 509.21 | 10 |
| 5 | 449.17 | 431.16 | 432.15 | 225.09 | F | 946.33 | 928.38 | 929.36 | 473.69 | 9 |
| 6 | 506.19 | 488.18 | 489.17 | 253.60 | G | 799.32 | 781.28 | 782.29 | 400.16 | 8 |
| 7 | 643.25 | 625.24 | 626.23 | 322.13 | H | 742.30 | 724.29 | 725.27 | 371.65 | 7 |
| 8 | 714.37 | 696.28 | 697.34 | 357.65 | A | 605.24 | 587.23 | 588.21 | 303.12 | 6 |
| 9 | 816.29 | 798.28 | 799.27 | 408.65 | C(-1.01) | 534.20 | 516.19 | 517.17 | 267.60 | 5 |
| 10 | 979.36 | 961.34 | 962.33 | 490.18 | Y | 432.20 | 414.19 | 415.17 | 216.60 | 4 |
| 11 | 1036.38 | 1018.36 | 1019.35 | 518.69 | G | 269.14 | 251.13 | 252.11 | 135.07 | 3 |
| 12 | 1093.40 | 1075.38 | 1076.38 | 547.20 | G | 212.11 | 194.10 | 195.09 | 106.56 | 2 |
| 13 | | | | | H(-.98) | 155.09 | 137.08 | 138.07 | 78.05 | 1 |

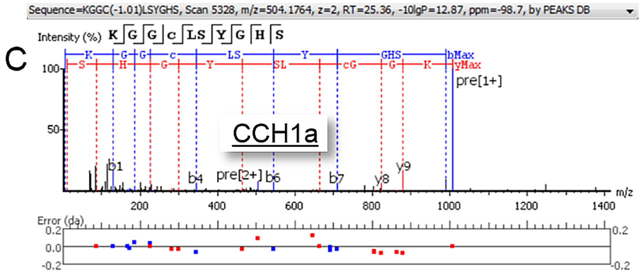


| # | b | b-H2O | b-NH3 | b (2+) | Seq | y | y-H2O | y-NH3 | y (2+) | # |
|---|---|---|---|---|---|---|---|---|---|---|
| 1 | 129.10 | 111.09 | 112.08 | 65.05 | K | | | | | 10 |
| 2 | 186.08 | 168.11 | 169.10 | 93.56 | G | 879.43 | 861.34 | 862.40 | 440.18 | 9 |
| 3 | 243.15 | 225.10 | 226.12 | 122.07 | G | 822.41 | 804.38 | 805.38 | 411.67 | 8 |
| 4 | 345.22 | 327.14 | 328.12 | 173.10 | C(-1.01) | 765.31 | 747.30 | 748.28 | 383.16 | 7 |
| 5 | 458.23 | 440.22 | 441.20 | 229.62 | L | 663.31 | 645.30 | 646.16 | 332.15 | 6 |
| 6 | 545.30 | 527.25 | 528.24 | 273.13 | S | 550.23 | 532.22 | 533.20 | 275.61 | 5 |
| 7 | 708.37 | 690.37 | 691.32 | 354.66 | Y | 463.23 | 445.18 | 446.17 | 232.10 | 4 |
| 8 | 765.35 | 747.34 | 748.32 | 383.17 | G | 300.17 | 282.12 | 283.14 | 150.57 | 3 |
| 9 | 902.41 | 884.40 | 885.38 | 451.70 | H | 243.11 | 225.10 | 226.08 | 122.05 | 2 |
| 10 | | | | | S | 106.05 | 88.04 | 89.02 | 53.52 | 1 |

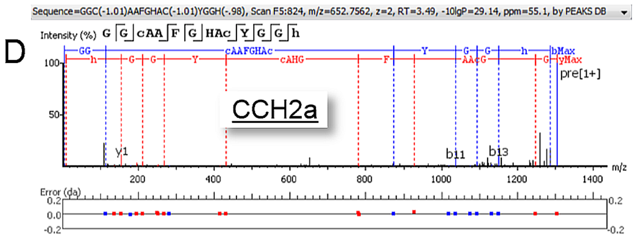


| # | b | b-H2O | b-NH3 | b (2+) | Seq | y | y-H2O | y-NH3 | y (2+) | # |
|---|---|---|---|---|---|---|---|---|---|---|
| 1 | 58.03 | 40.02 | 41.00 | 29.51 | G | | | | | 14 |
| 2 | 115.05 | 97.04 | 98.02 | 58.03 | G | 1247.48 | 1229.47 | 1230.46 | 624.24 | 13 |
| 3 | 217.05 | 199.04 | 200.03 | 109.03 | C(-1.01) | 1190.46 | 1172.45 | 1173.43 | 595.73 | 12 |
| 4 | 288.09 | 270.08 | 271.06 | 144.54 | A | 1088.46 | 1070.45 | 1071.43 | 544.73 | 11 |
| 5 | 359.13 | 341.12 | 342.10 | 180.08 | A | 1017.42 | 999.41 | 1000.40 | 509.21 | 10 |
| 6 | 506.19 | 488.18 | 489.17 | 253.60 | F | 946.39 | 928.35 | 929.36 | 473.69 | 9 |
| 7 | 563.22 | 545.21 | 546.19 | 282.12 | G | 799.32 | 781.29 | 782.31 | 400.16 | 8 |
| 8 | 700.28 | 682.26 | 683.25 | 350.64 | H | 742.30 | 724.29 | 725.27 | 371.65 | 7 |
| 9 | 771.31 | 753.30 | 754.29 | 386.16 | A | 605.24 | 587.23 | 588.21 | 303.12 | 6 |
| 10 | 873.31 | 855.30 | 856.29 | 437.16 | C(-1.01) | 534.20 | 516.19 | 517.17 | 267.60 | 5 |
| 11 | 1036.38 | 1018.36 | 1019.35 | 518.69 | Y | 432.20 | 414.19 | 415.17 | 216.60 | 4 |
| 12 | 1093.40 | 1075.39 | 1076.38 | 547.20 | G | 269.14 | 251.10 | 252.11 | 135.07 | 3 |
| 13 | 1150.42 | 1132.41 | 1133.39 | 575.71 | G | 212.11 | 194.10 | 195.09 | 106.56 | 2 |
| 14 | | | | | H(-.98) | 155.09 | 137.08 | 138.07 | 78.05 | 1 |

**Figure S3. Quadrupole orbitrap MS$^2$ spectra generated by ESI-Q Exactive Orbitrap MS brain and midgut extracts of *Ae. aegypti* males for sequence identification of CCHamides.** Resulting fragmentation spectra revealed the presence of CCHa1 and CCHa2 in the brain (A, B) and in midgut extracts (C, D) of male adult mosquitoes.
